# Vision Transformers as a Framework for Imaging Fluorescence Correlation Spectroscopy

**DOI:** 10.64898/2026.10.21.760315

**Authors:** Shao Ren Sim, Zijie Wang, Shambhavi Pandey, Shiyue Zhou, Thorsten Wohland

## Abstract

Fluorescence correlation spectroscopy (FCS) is widely used for quantifying molecular dynamics in live cells. However, conventional FCS relies on the autocorrelation function (ACF), a lossy transformation that discards higher-order information, mathematically couples distinct biophysical parameters, and requires long measurements (*∼*50000 frames, or *∼* 1 minute) that obscure short-lived transient events. To overcome this information bottleneck, we introduce ImFCSViT, a spatiotemporal Vision Transformer-based framework that bypasses the ACF representation entirely, operating directly on raw fluorescence image stacks. By introducing a dedicated normalization token to preserve global intensity statistics, our approach simultaneously extracts four biophysical parameters (diffusion coefficient *D*, particle density *N*, particle brightness *B*, and PSF structure factor *σ*_0_) in a single forward pass. Requiring only 2.5 s of data, and potentially down to 1 s (a 1-2 order reduction in data requirement), our approach accurately recovers diffusion coefficients while utilizing its spatial field-of-view to reveal spatially heterogeneous, slow-diffusing sub-populations in lipid bilayers and live-cell plasma membranes that are invisible to classical FCS. By leveraging the data-driven representational learning capabilities of deep learning, we present a flexible, generalizable, and powerful tool for mapping complex, multi-parameter spatiotemporal dynamics in biological systems.

**SIGNIFICANCE**
Standard fluorescence correlation spectroscopy (FCS) is severely bottlenecked by long measurement times and lossy mathematical approximations, making it difficult to observe fast cellular events or decouple interrelated physical properties. To address these limitations, we introduce the ImFCSViT framework, based on a Vision Transformer that bypasses the lossy data averaging of conventional methods, operating directly on raw microscopy videos. By capturing both spatial context and global intensity statistics, our model estimates four key biophysical properties with total measurement times as low as one second, making it compatible with standard imaging approaches, and uncovering spatially heterogeneous structural domains in live cell membranes. This approach provides researchers with a fast, flexible tool to observe biological dynamics that are inaccessible to classical approaches.

## INTRODUCTION

Since its initial introduction, fluorescence correlation spectroscopy (FCS) has been a mainstay in the analysis of molecular dynamics, enabling the extraction of critical biophysical parameters such as the diffusion coefficient (D) and apparent particle density (N) within an observation volume. The foundation of FCS is the autocorrelation function (ACF), which computes the similarity of intensity fluctuations against a time-shifted variant over multiple lag times (1).

Despite its widespread use, the ACF itself comes with its own limitations. Conventional FCS fitting relies on mechanistic fit functions, which use approximations to encode our understandings and interpretations of the underlying physics into a mathematical descriptor. However, the ACF itself only considers correlations up to the second order, disregarding higher-order kinetic processes that also contribute to the recorded intensity fluctuations. Furthermore, ACFs are biased estimators that require sufficient statistics for accurate parameter estimation, which in practice, necessitates a minimum measurement time of 50000 frames (∼ ∼50 seconds at a 1.0 ms frame time) or more (2, 3), restricting the utility of FCS in tracking transient biophysical phenomena.

In an attempt to modernize FCS analysis, deep learning has increasingly been adopted in FCS-based workflows. Several recent approaches have successfully utilized neural networks to replace or augment traditional non-linear least squares (NLS) fitting, offering tools for classifying anomalous diffusion modes (4), artifact segmentation (5), and accelerating parameter extraction (6).

While these models represent significant advancements, they are still reliant on the ACF, and thus unable to utilize the higher-order information discarded in the correlation process. To fully leverage the capacity of deep learning on high-dimensional datasets, recent methods that bypassed the ACF representation entirely have been proposed, operating directly on raw recorded photon counts. These approaches shift the focus from functional approximation (*i*.*e*., replicating the fit function used for data labeling) to representational learning (*i*.*e*., extracting the optimal features directly from the raw data). In essence, this approach does not assume that the ACF is the ideal representation, and instead dynamically derives the feature representation that is most optimized for the task at hand.

One of the main pioneering works in this regard is ImFCSNet (7), a convolutional neural network (CNN)-based approach that directly operated on raw image stacks. This method dramatically increased temporal resolution, allowing for diffusion coefficient estimates in just 2500 frames, a 20*×* reduction compared to conventional FCS fitting. Subsequent work then extended ImFCSNet to predict particle densities, extracting insights that would have been inaccessible or smoothed over by the averaging effect of correlations (8). In concert, both these approaches show that deep learning is not only feasible with raw photon counts, but capable of uncovering representations free from the lossy transformation to the ACF.

While ImFCSNet has overcome the temporal bottleneck of standard FCS, it is, by design, only able to predict individual parameters in isolation, requiring bespoke setups for each target. Considering physical properties in isolation fundamentally ignores the fact that these biophysical properties are coupled. For example, standard FCS mathematical models derive an apparent particle number (*N*), a confounding term that combines true particle density with individual particle brightness (9). Similarly, the diffusion coefficient (*D*) and the point-spread-function (PSF) are mathematically linked. Estimating each property in isolation fails to account for cross-parameter covariances. Furthermore, ImFCSNet is itself a custom built CNN architecture, limiting interpretability and hindering cross-pollination across disciplines, where advances in deep learning could be translated into physical insights that help further our understanding with analytical fitting.

To bridge this gap, we build upon the foundations set out by ImFCSNet, and design ImFCSViT, a deep learning framework that handles multi-parameter prediction. To this end, we adopt the standard Vision Transformer (ViT) architecture (10). Unlike CNNs, whose strengths lie in the extraction of local overlapping spatial patterns, transformer-based architectures are suited for sequence-based learning through the self-attention mechanism, and well equipped to extract continuous temporal features across long time-horizons, making them ideal for fluorescence fluctuation traces. Furthermore, using a well-understood standard means we can also take advantage of insights and advancements made available through the deep learning field as a whole, allowing us to build up interpretability without needing to build up intuition from scratch.

We follow the simulation-driven training paradigm established by (7) and (8), generating synthetic photon-count traces to provide decoupled ground-truth labels for all four parameters. By generating synthetic photon-count traces from simulated diffusion and optics models, we embed the relevant physics directly into the training data rather than into an approximate fit function. This shifts the role of theory from analytic approximation to generative prior, and it is currently the only feasible route to obtaining a comprehensive, noise-diverse dataset needed to train a robust multi-output model.

Table 1 summarizes the predictive capabilities and temporal requirements of our proposed ImFCSViT framework against existing analytical and deep learning methods.

**Table 1:** Comparison of parameter estimation capabilities and required measurement times across analytical and deep learning methods.(^∗^ Note: While ImFCSViT successfully extracts relative brightness scaling, calculating absolute quantitative stoichiometry currently requires further noise-aware training; see Discussion and Appendix A.2.)

| Method | Diffusion<br>($D$) | Particle Density<br>($N$) | Brightness<br>($B$) | PSF Calib.<br>($\sigma_0$) | Req.<br>Time |
| --- | --- | --- | --- | --- | --- |
| Conventional FCS | ✓ | ✓ | × | × | 50 s |
| Number & Brightness | × | ✓ | ✓ | × | 20 s |
| ImFCSNet (CNN) | ✓ | × | × | × | 2.5 s |
| **ImFCSViT (ViT)** | ✓ | ✓ | ✓* | ✓ | **2.5 s** |

**Table 2:** Computational complexity and runtime benchmarking for a 128 *×* 128 *×* 2500 frame data chunk. MFLOPs are reported per single model forward pass (i.e., per 3×3 ROI for ImFCSNet and the 3 *×* 3 ImFCSViT, and per pixel for 1 *×* 1 ViT). Total runtimes represent the total inference time for the full spatial grid. Speedups are normalized against the ImFCSNet baseline.

| Model | Params | MFLOPs | Runtime (s) |  | Relative Speedup |  |
| --- | --- | --- | --- | --- | --- | --- |
|  |  |  | CPU | GPU | CPU | GPU |
| ImFCSNet ($3 \times 3$) | 715276 | 307.3 | 44.31 | 4.91 | 1.00× | 1.00× |
| ImFCSViT ($3 \times 3$) | 592420 | 119.6 | 44.43 | 5.36 | 0.99× | 0.91× |
| ImFCSViT ($1 \times 1$) | 39748 | 3.9 | 8.05 | 1.41 | 5.50× | 3.48× |

In this work, we demonstrate that the ImFCSViT framework is able to disentangle multiple coupled physical parameters simultaneously, generalizing across the simulation-to-experimental data domain gap. We then extend our validation to cells, showing that our generalized training procedures can be easily adapted to alternative systems without loss of fidelity. Finally, we highlight key limitations of simulation-led model training by showcasing failure cases with complex physical morphologies that are not represented in our training data, and utilize the inherent flexibility of ViT architectures to address these concerns with mitigation strategies.

## MATERIALS AND METHODS

### Imaging FCS

Historically, classical FCS techniques generally revolved around single point measurements, generally acquired using confocal microscopy systems (1, 11±13). While modified approaches existed to extend this analysis into 2 dimensions (14±17), these approaches were limited in terms of spatial resolution, and could not provide a contiguous view of the sample under measurement.

With the introduction of area-based detectors such as electron multiplying charge-coupled devices (EMCCDs), scientific complementary metal-oxide-semiconductor (sCMOS) or quantitative CMOS (qCMOS) cameras, and single-photon avalanche diodes (SPAD) arrays alongside planar illumination techniques, it became possible to perform multiplexed FCS analysis. This approach, known as Imaging FCS, was explicitly designed to incorporate both temporal and contiguous spatial information (18).

The basis of FCS analysis is the fact that the fluorescent fluctuations around equilibrium contain information about the particle dynamics within the observation volume (19). To extract said information, the autocorrelation function (ACF) is used, where the self-similarity of a signal is computed by correlating a fluorescence signal *F* (*t*) against itself, but delayed by a certain time *τ* (20).

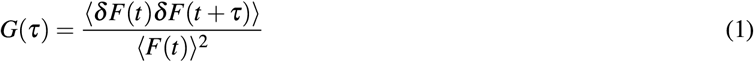

The angled brackets denote time averaging over the total measurement time 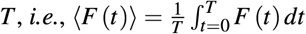.

As the ACF encodes the probability of a molecule entering and leaving the illuminated observation region, mathematical fit models can be derived to extract parameters of interest through non-linear least squares fitting. Conventionally, Imaging FCS fitting uses the iterative Levenberg-Marquardt algorithm to fit these models (21, 22).

In this work, we focus on the simple case of 2-dimensional free diffusion of a single molecular species under total internal reflection fluorescence (TIRF) illumination (23). Here, the mechanistic fit model aims to fit for the diffusion coefficient *D*, and the apparent particle concentration per-pixel *N*, taking the following form.

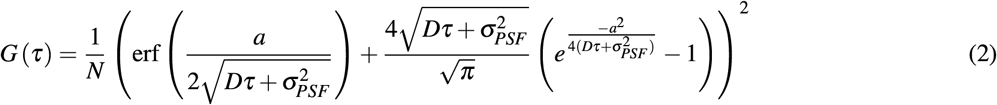

where *a* is the length of a square observation area in object space, *σPSF* is the PSF width (*i*.*e*., the standard deviation of the Gaussian illumination profile) given by *σPSF* = *σ*_0_*λ*_*em*_*/NA* where *λ*_*em*_ is the emission wavelength, *NA* is the numerical aperture, and *σ*_0_ is the structure factor of the point-spread function (PSF) determined through calibrations.

By repeating this procedure across all available pixels acquired from the area-based detector, we can obtain parameter maps, where the spatial information made available through the multiplexed FCS setup makes it possible to visualize how particle dynamics change over space.

Crucially, Equation 2 highlights the inherent mathematical coupling of these biophysical parameters. The temporal descriptor *D* and the spatial descriptor *σ* are linked as a sum 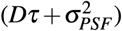, meaning models are unable to fit both simultaneously. In practice, this is handled by using a fixed PSF *σ*_*PSF*_ term as determined through calibrations, but this also means the fit function is unable to account for PSF shifts over time, such as microscope defocusing or calibration drift, inevitably corrupting the *D* estimates.

Furthermore, Equation 2 does not account for molecular brightness *B*, only providing an estimate of the apparent number of particles *N*. While it is possible to derive brightness naively through *B* = ⟨*F* ⟩*/N*, this estimate is biased due to the presence of background contributions (24). In general, decoupling particle densities from particle brightness requires more specialized techniques like number and brightness (N&B) analysis.

Finally, because correlation functions are fundamentally biased estimators, a sufficiently long measurement time is computationally required to obtain robust fits, generally necessitating 50 000 frames (or 50 seconds with a 1.06 ms frame time) as a standard (3). While it is possible to work around this limitation through small chunk analysis (25) or fit function formulations that accounts for shorter measurement times (26), this still limits the temporal resolution of analysis, which potentially obfuscates any physical changes that may occur at a faster rate.

### Number and Brightness Analysis

As standard Imaging FCS conflates the true particle density with molecular brightness into the apparent *N* term, an alternate mathematical approach is required to accurately uncouple these variables. Number and brightness (N&B) analysis is a statistical method that allows for the determination of apparent brightness over an image (27). N&B analysis was first proposed in (28, 29), and demonstrated on live cell measurements in (9). N&B analysis can similarly map multiplexed predictions using TIRF illumination and EMCCD detectors (24).

The intensity value at any given point in time, *I*(*t*), is a function of the number of particles *N*(*t*) and the brightness of each particle *B*. To decouple these parameters, we analyze the time-averaged intensity ⟨*I*⟩, described in Equation 3.

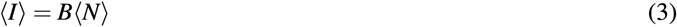

To independently determine the number *N* and brightness *B* of particles in a given measurement, N&B utilizes the raw statistical moments of the sample’s time-varying intensity profile (29, 30). For an EMCCD camera, the apparent number *N* and apparent brightness *B* of particles in the observation volume can be defined as

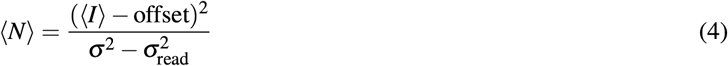

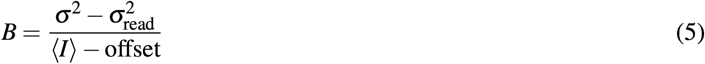

where offset is the intensity offset characteristic to the acquisition setup, *σ* ^2^ is the variance of the signal, and 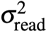 is the readout noise variance of the camera.

Because it explicitly utilizes statistical variance, N&B analysis is highly sensitive to background noise artifacts including dark counts, stray light, autofluorescence, and immobile fractions (27). While sufficiently large signals mean the background is negligible, background correction is generally required. This is achieved by subtracting the background offset (measured without sample illumination) and accounting for systematic noise by subtracting 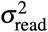 (calculated directly from the covariance of a dark image stack) (31).

Crucially, while FCS and N&B analysis operate on the exact same raw fluorescence data, they utilize different mathematical paradigms, and require different measurement conditions. As such, despite the possibility of determining both mobility (*D*) and spatial (*N, B* and the PSF) descriptors simultaneously (32), it is not yet possible to do so in a single unified framework. In this work, we specifically seek to leverage the representation learning capabilities of deep learning to derive both kinematic and spatial parameters with a single model.

### Vision Transformers

In this work, we transition away from convolutional deep learning pipelines, and instead utilize the Vision Transformer (ViT) architecture from (10). While initially designed for natural language processing (33), transformers are a general architecture that is agnostic to the input modality due to the minimal inductive biases present in the architecture. It has been successfully applied in static vision (34) and videos (35±37), making it uniquely suited for our input modality of highly correlated, spatiotemporal fluorescence image stacks.

The full model schematic used in this work is shown in Figure 1. Following the video tokenization strategy from (38) and (35), we parse the input by splitting the input image stacks into non-overlapping spatiotemporal tubelets. Taking inspiration from the spatial constraints of ImFCSNet (7), we process a total input horizon of 2500 frames, but chunk the data into localized 3 × 3 × 10 tubelets (representing *W* × *H* × *T*). Each tubelet is then embedded into a 1-dimensional vector representation of length 96. Through this operation, any given 2500-frame sequence is natively embedded into a token sequence of length 250. While this specific setup is our canonical configuration, we can adapt to different input sizes or different frame counts as needed, which we show in the Results and Appendix D.2.

**Figure 1:**
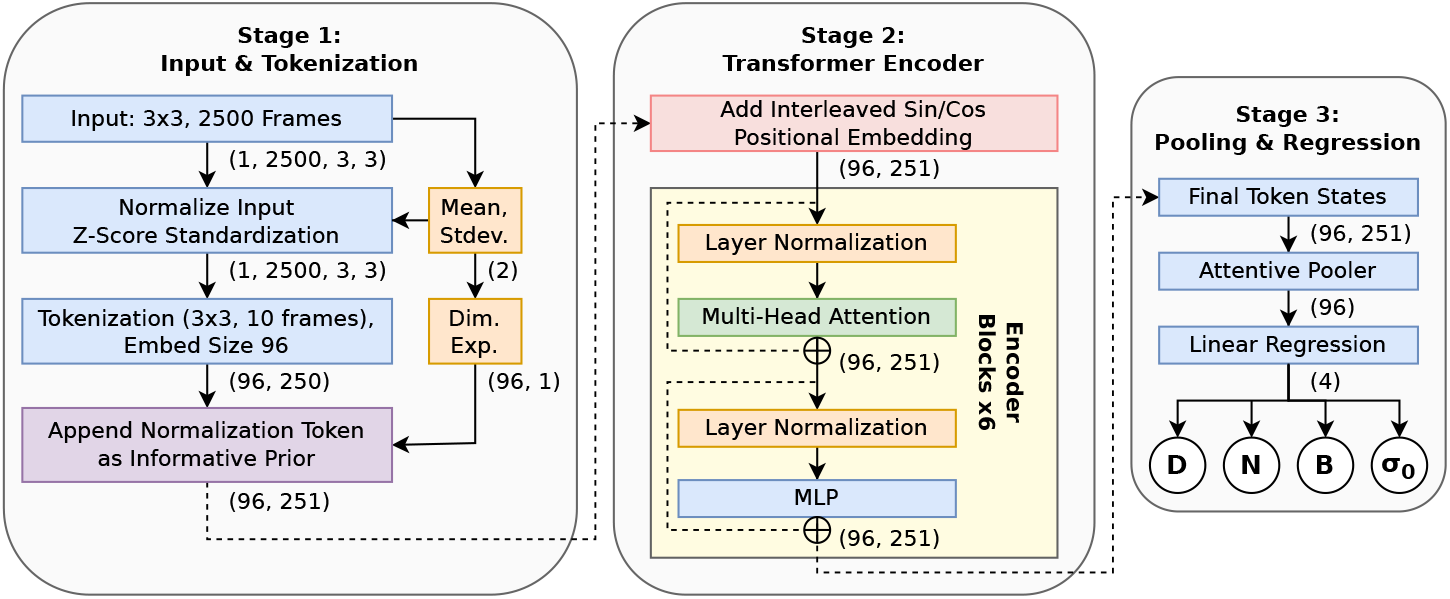
Schematic overview of the proposed spatiotemporal ImFCSViT architecture, based on the standard Vision Transformer (ViT). The pipeline is divided structurally into three stages. **Stage 1 (Input & Tokenization)** illustrates the dual-path preprocessing, where the raw image stack is normalized through Z-score standardization without discarding the bulk statistics (mean and standard deviation). To make the statistics accessible to the subsequent transformer blocks, they are dimensionally expanded through linear projection into a dedicated normalization token that gets appended to the sequence. **Stage 2 (Transformer Encoder)** processes the 251-token sequence through 6 consecutive transformer encoder blocks. **Stage 3 (Pooling & Regression)** aggregates the final token states into a single unified feature vector of length 96, which is then mapped through a linear regression head to simultaneously predict the four parameters of interest (*D, N, B*, and *σ*_0_).

Crucially, standard ViT implementations are insufficient for the simultaneous prediction of coupled biophysical parameters. Given an unprocessed fluorescence intensity trace, the most dominant features are the mean and standard deviation. These features are important for the determination of particle densities (*N*) and brightness (*B*) (9, 27), but they are tangential to the inter-frame fluctuations that encode the movement of particles, becoming a nuisance variable for diffusion coefficient (*D*) fitting. Previous deep learning approaches were forced to choose: (7) achieved robust diffusion estimates by applying Z-score standardization, effectively destroying the bulk statistics needed for *N* and *B*. Conversely, (8) utilized bespoke task-specific normalizations, requiring separate pipelines for different targets. Neither method is ideal for our use case of simultaneous multi-parameter fitting.

To resolve the conflicting requirements of our different targets, the core modification we introduce in our ImFCSViT variation is the inclusion of a dedicated “normalization token”. As detailed in Equation 6, we apply Z-score normalization to each pixel *x* to isolate the kinetic fluctuations necessary for diffusion tracking:

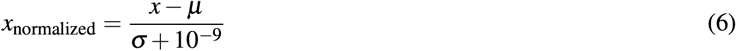

where *µ* and *σ* are the mean and standard deviation computed over all pixels and frames of the 3 *×* 3 *×* 2500 input tensor (*i*.*e*., per-sample global statistics). A numerical stability term is included in the denominator to guard against zero-division. Unlike conventional normalization, we do not discard the statistics, instead using a linear projection to cast *µ* and *σ* directly into a length-96vector, matching the dimensions of the spatiotemporal patch tokens, and thus preserving the global intensity context without corrupting the kinetic fluctuations.

This normalization token is then appended to the input sequence via an early-fusion configuration (39). By supplying this token at the onset, the ImFCSViT model can simultaneously process the normalized input for optimal *D* fitting, while also having access to the global context required for *N* and *B* estimation. Ablation studies (Appendix F) confirm this, as attempting multi-parameter fitting on unnormalized data results in diffusion estimates (*D*) that suffer a drop in accuracy and consistency, whereas standard normalization without our proposed token fundamentally prevents the accurate extraction of particle density (*N*) and brightness (*B*).

To encode the token ordering into the sequence, we use interleaved sin/cos positional embeddings as per (40). The token-space representations from the terminal transformer block are then passed through an attentive pooler (38, 41), acting as a non-linear feature aggregation layer that reduces the sequence into a unified feature vector. Finally, multivariate linear regression is used to learn a mapping from the 96-length feature vector to our 4 target parameters.

### Model Training

Our model is optimized via fully supervised learning, computing the overall training loss using a standard mean squared error (MSE) metric across all simultaneous target predictions. As experimental datasets lack ground-truth labels for our parameters of interest, we rely entirely on physical simulations to generate input-target pairs.

One critical design decision of our simulation pipeline is the use of wide parameter variations during training. The practical consideration is that deep learning regression models generally fail to extrapolate beyond the bounds of their training distribution. To ensure generalizability and robustness, we deliberately sample across an expansive physical parameter space during simulation, training the network on bounds beyond expected experimental ranges (e.g., simulating extreme diffusion regimes rather than exclusively expected slow variants). These parameters are sampled log-uniformly, ensuring that accuracy and precision can be optimized for across all parameter scales.

The data simulation process is detailed in Appendix C.1. In short, we use simple, 2-dimensional single-component free diffusion simulations as per (8). Instead of relying on simulation complexity, we adopt a regularization-heavy approach to bridge the simulation-to-experiment gap. We augment the training sets with simulated EMCCD noise profiles and low signal-to-noise ratio (SNR) conditions, forcing the network to learn feature representations robust to noise.

Because our models simultaneously predict physical parameters that vary significantly in scale and across multiple orders of magnitude, applying a raw MSE loss would structurally bias the network toward minimizing larger-magnitude parameters. To prevent this, we apply a logarithmic min-max normalization to the targets using the known absolute upper and lower bounds of our simulation space (Equation 7) prior to the MSE calculation, ensuring that the loss gradients contributed by each target are scale-invariant and bounded within the range [0, 1].

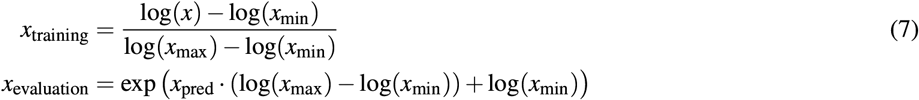

During inference, predictions are reverted to physical units via the inverse transformation. Detailed training configurations and hyperparameter settings are provided in Appendix C.

Because our model minimizes MSE in log-space, its predictive errors are symmetric in the logarithmic domain, which manifests as multiplicative (rather than additive) errors in linear space. Furthermore, our parameters of interest (namely *D, N*, and *B*) are strictly positive and typically vary across multiple orders of magnitude in biological samples, making the geometric mean an inherently more robust measure of central tendency.

Crucially, the arithmetic mean does not commute with non-linear transformations due to Jensen’s inequality (42). Because the exponential function is strictly convex, the arithmetic mean of exponentiated predictions is always strictly greater than the exponentiated arithmetic mean of log-space predictions (*E* [*e*^*x*^] ≥ *e*^[*x*^). Exponentiating the arithmetic mean of our model’s log-space predictions yields the geometric mean in linear units. To avoid transformation-induced bias and to ensure our summary statistics are directly comparable regardless of the underlying estimator’s loss function, we report geometric means for all methods (both ImFCSViT and NLS). Correspondingly, variation is also reported using the geometric standard deviation factor, denoted as the multiplicative interval [*µ*_g_ */σ*_g_, *µ*_g_ *× σ*_g_], which represents *±*1 standard deviation in log-space. The exception is the PSF *σ*_0_ parameter, which is predicted and optimized in linear space without a logarithmic transform, and is therefore reported using the arithmetic mean and standard deviation.

This overall framework also allows our core architecture and training procedures to remain fixed across different applications. All reported ImFCSViT models in this work share identical training and simulation hyperparameters, with the only differences being the simulation settings related to the experimental setup, namely: the fluorophore-specific emission wavelength, camera-specific dark noise pattern, and sample-specific PSF calibrations. This provides confidence that our proposed framework is generalizable to diverse experimental settings entirely through physical prior modification, eliminating the need for secondary end-user hyperparameter tuning.

## RESULTS

To verify that the ImFCSViT framework is robust across a wide range of biological systems and experimental conditions, we apply our approach on a variety of different biophysical experiments, and compare the model predictions against conventional non-linear least squares (NLS) fitting where applicable.

It should be emphasized that the two approaches probe different temporal and spatial scales. Standard NLS fitting extracts bulk dynamical parameters averaged across a full 50 000-frame observation window (50 seconds assuming a 1 ms frame time). In contrast, our ImFCSViT framework, by design, evaluates a much shorter 2500-frame horizon (2.5 seconds). Because of this 20-fold difference in temporal resolution, parameter estimates in non-stationary biological systems will naturally differ. NLS provides a long-term, time-averaged metric, whereas ImFCSViT provides 20 localized estimates in the same given window. As such, parity between the two methods is not expected in heterogeneous samples where dynamics shift over time. Furthermore, ImFCSViT model takes in a localized 3 *×* 3 pixel region of interest (ROI). Given our simulations used for training, our model strictly assumes spatial homogeneity across all 9 pixels within this region (*i*.*e*., all particles in our simulated region share the same diffusion coefficient, fluorescent particles are distributed evenly etc.).

All samples were pre-processed with pixel-wise polynomial trend correction (order 4) to mitigate photobleaching. For all analyses, we employed a sliding window with a spatial stride of 1 (overlap mode as described in (7)), which provides higher-resolution parameter maps at the cost of spatial correlation between adjacent predictions.

Detailed experimental procedures and sample preparations are provided in Appendix B.

### DOPC Bilayers Under Variable Concentrations and Illumination Power

To validate the model experimentally, we imaged DOPC supported lipid bilayers doped with the rhodamine-labelled lipid RhoPE at various molar concentrations and under multiple illumination powers to assess all four ImFCSViT outputs simultaneously (Appendix B.1).

As shown in Figure 2a, a naive comparison of global ImFCSViT estimates against NLS-derived diffusion coefficients (*D*) initially suggests the ImFCSViT-derived *D* values are systematically biased across all available conditions by 20 − 40%, suggesting a systemic discrepancy. However, this comparison is inappropriate. Inspecting the distribution of *D*-values reveals a long tail of slow outliers (Figure 2b). Inspecting these low-*D* regions spatially show that they map to high brightness regions, corresponding to slow-moving or immobile residual vesicles, an artifact of the supported lipid bilayer forming process (Figure 2c). These slow-moving particles are not captured by conventional 1-component FCS fitting. Because long-horizon temporal correlation is dominated by the high-frequency fluctuations of the fast-moving majority, the low-frequency signatures of hindered particles act as a minority component that is indistinguishable from baseline noise. In contrast, ImFCSViT evaluates 3 *×* 3 spatial windows, allowing it to distinguish hindered, quasi-stationary particles from pure stochastic noise. Consequently, ImFCSViT manages to extract both populations from the raw intensity traces.

**Figure 2:**
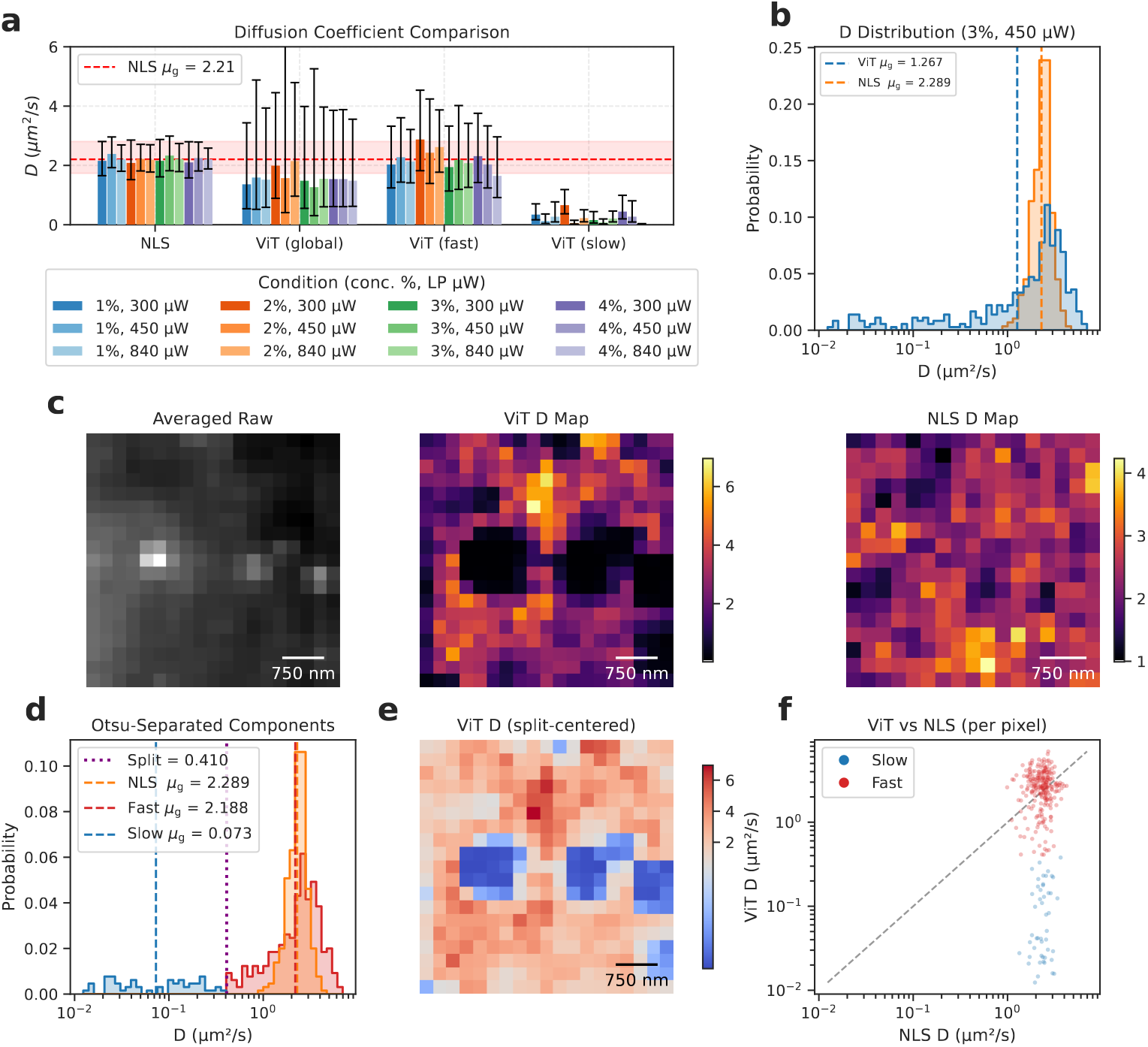
Validation of ImFCSViT diffusion analysis on DOPC supported lipid bilayers. (**a**) Diffusion coefficients (*D*) for NLS (control) and ViT measurements. For each ViT sample, three values are shown: the global (naive) *D* from all trajectories, and the *D* of the fast and slow sub-populations separated by automatic Otsu thresholding. Global *D* systematically underestimates the true value; the fast component recovers the NLS reference value. Bars represent geometric mean *×/÷* geometric standard-deviation factor (*i*.*e*., the interval [*µ*_g_*/σ*_g_, *µ*_g_ *× σ*_g_]) for all methods. The red horizontal band marks the geometric mean and geometric SD interval of the pooled NLS measurements. (**b**) Log-scale distribution of *D* for a representative measurement (3% concentration, 840 µW laser power), revealing a long tail of slow outlier values. Vertical lines mark the geometric means of the ViT global (blue) and NLS (orange) distributions. (**c**) Temporally averaged 2500-frame region and the corresponding ViT and NLS *D* map, showing that slow-*D* regions in the ViT map coincide with high-intensity residual vesicles, but are absent in NLS. (**d**) Otsu-separated fast and slow components of the same measurement as (b), with the automatic split point (purple) and the geometric means of the NLS (orange), fast (red), and slow (blue) populations. (**e**) *D* map with a diverging colour scale centred on the Otsu threshold, highlighting fast (red) and slow (blue) pixels that correlate with visible vesicles in the raw image. (**f**) Scatter plot of per-pixel *D* values of the ViT versus the NLS *D* values, showing that the ViT-derived fast-component at 2500 frames correlates well with the NLS estimations at 50 000 frames.

To isolate the bulk diffusion coefficient, we apply Otsu’s algorithm to the log-space *D* distribution, automatically identifying a threshold that separates the fast and slow components by maximizing inter-class variance (43). Using this method, the fast-*D* component matches the NLS-derived *D* values (Figure 2d), confirming that ImFCSVIT correctly estimates the bilayer diffusion when vesicle contamination is accounted for. Conversely, the slow-*D* component always co-localizes with visible vesicles in the raw intensity images (Figure 2e), indicating that the variations observed by the ViT model are not spurious uncertainties, but true dynamic heterogeneity that is otherwise obscured by the assumptions and approximations present in FCS-based fitting, being indistinguishable from background noise to the temporal correlation process. In contrast, ImFCSViT, operating on raw intensity traces, manages to extract both components. Furthermore, we observe that the existence of slow and fast components are consistent across all 12 samples (Appendix A.3).

Beyond molecular mobility, we evaluate the model’s ability to estimate intensity-derived parameters. As shown in Figure 3a and 3b, ImFCSViT provides consistent particle density estimates (*N*) for a given molecular concentration independent of increasing laser powers, and vice versa with brightness estimates (*B*). This is in contrast to conventional NLS Fitting, where the estimated *N* values decrease with increasing laser power (caused by the increase in the signal-to-noise ratio and corresponding inverse relationship to *N* as per Equation 2). Notably, ImFCSViT estimates also scale proportionally with the known physical changes in concentration and laser power, demonstrating that our simulation-led deep learning framework is able to decouple concentration from particle brightness. Our model also achieves this without requiring a background correction step as per N&B analysis, though the lack of explicit background modelling does mean ImFCSViT is unable to handle situations where quantitative accuracy of intensity-derived parameters are required (we examine the applicability of the model’s CPS output in place of N&B analysis for stoichiometry in A.2).

**Figure 3:**
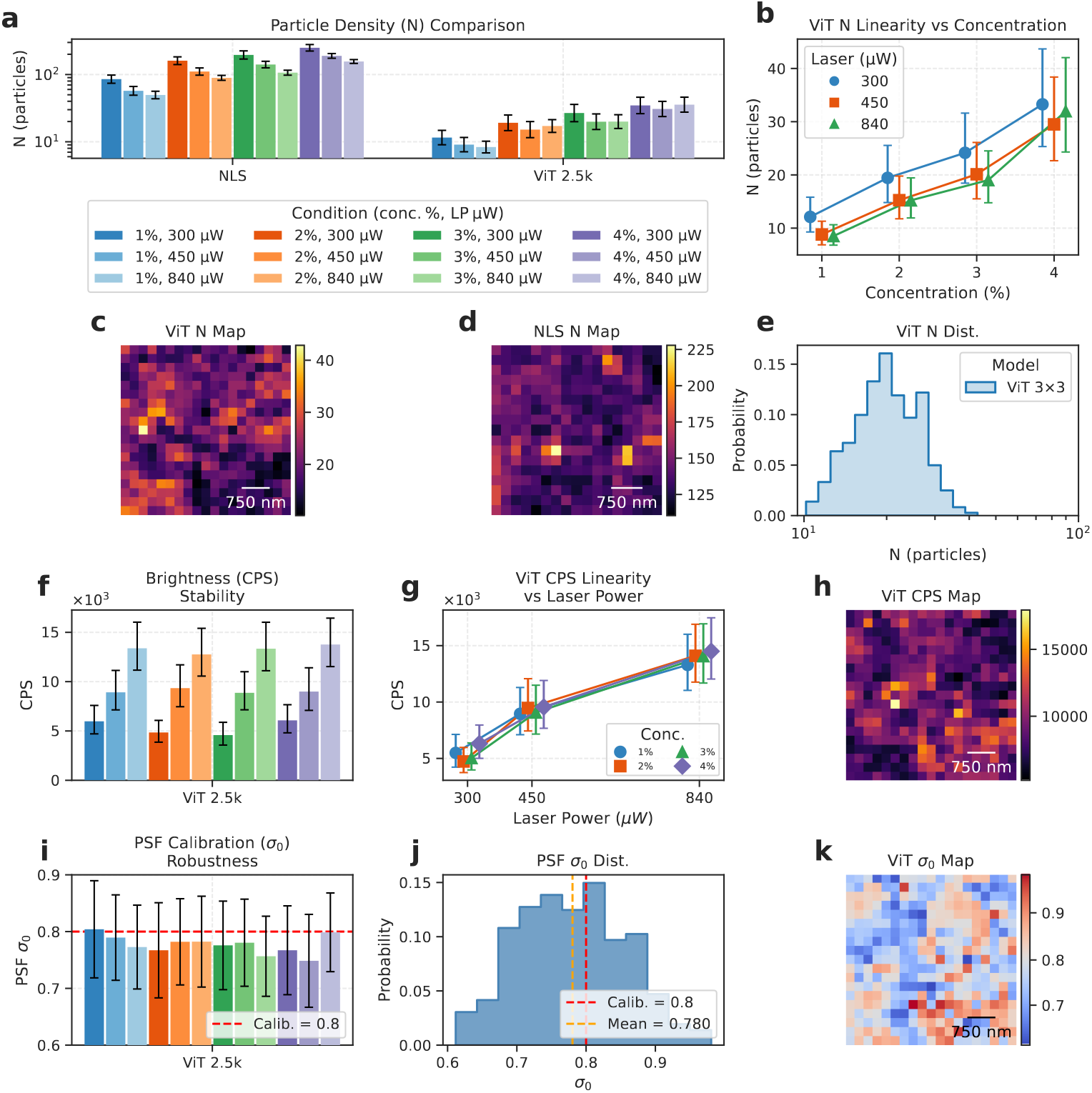
Validation of auxiliary ImFCSVIT outputs on DOPC supported lipid bilayers. (**a**) Particle density (*N*) for NLS (control) and ViT measurements. NLS predicted N values decrease with increasing laser power, whereas ViT N estimates remain relatively constant. Bars represent geometric mean *×/÷* geometric standard-deviation factor for all methods. (**b**) ViT *N* as a function of label concentration, coloured by laser power, demonstrating a linear response and relative consistency irrespective of laser power. (c±e) Spatial maps of ViT *N* (**c**) and NLS *N* (**d**) for a representative measurement from Figure 2c (3% concentration, 450 µW laser power), and the log-scale distribution of per-pixel ViT *N* values (**e**). (**f**) Brightness (CPS) across ViT measurements, showing stability against sample condition and laser power. (**g**) ViT CPS as a function of laser power, coloured by label concentration, showing proportional scaling with illumination laser power. (**h**) Spatial map of ViT CPS for the same representative measurement. (**i**) PSF width (*σ*_0_) across ViT measurements, with the calibrated reference value of 0.8 (red dashed line), demonstrating consistency across all experimental conditions. Bars represent arithmetic mean *±* SD. (**j**) Linear-scale distribution of per-pixel ViT *σ*_0_ values for the representative measurement, centred around the calibration target. (**k**) Spatial map of *σ*_0_ with a diverging colour scale centred at 0.8, highlighting local deviations from the calibrated PSF width which is not accessible via standard PSF calibration methods.

Finally, we analyzed the network’s predictive mapping of the point-spread-function (PSF) calibration factor (*σ*_0_). While the true optical PSF remained stable during these specific *in vitro* measurements, the network accurately and consistently predicted the known calibration baseline (Figure 2i). ImFCSViT also derives the PSF estimate on a 3 *×* 3 ROI basis, which is in contrast to the standard calibration approach, which only provides a singular global constant across a given image. Furthermore, we show in the Appendix A.1 that our model is capable of quantifying the PSF changes across an induced defocusing event, all while retaining the consistent *D* value that is robust to defocusing as demonstrated in ImFCSNet in (7).

### Live-Cell Dynamics

Having validated the approach on controlled DOPC bilayers, we next applied the ImFCSViT framework on live-cell measurements. We imaged CHO-K1 cells transiently expressing PMT-mApple, a plasma-membrane-targeted fluorescent protein serving as a freely diffusing membrane probe (Appendix B.2). Background pixels were excluded by automated Otsu thresholding applied to the time-averaged raw image.

A naive analysis of the global ImFCSViT *D* distribution again yields a geometric mean (0.22 and 0.21 *µm*^2^*/s* for Cell 1 and Cell 2) that lies well below the NLS-derived estimates (0.74 and 1.01 *µm*^2^*/s*). However, the log-scale histograms of per-pixel ImFCSViT *D* values (Figure 4c,g) reveal a clearly bimodal distribution, with the dominant slower mode accounting for the downward bias of the global statistic. Separating the two populations through non-parametric Otsu classification recovers a fast component whose geometric mean (0.99 and 0.95 *µm*^2^*/s*) occupies a region consistent with NLS estimates and their respective uncertainties, and a slow component comprising 40-45% of pixels in both cells, populating a region largely absent from the NLS D-distribution, and suggesting that membrane organisation or probe behaviour is more heterogeneous than anticipated.

**Figure 4:**
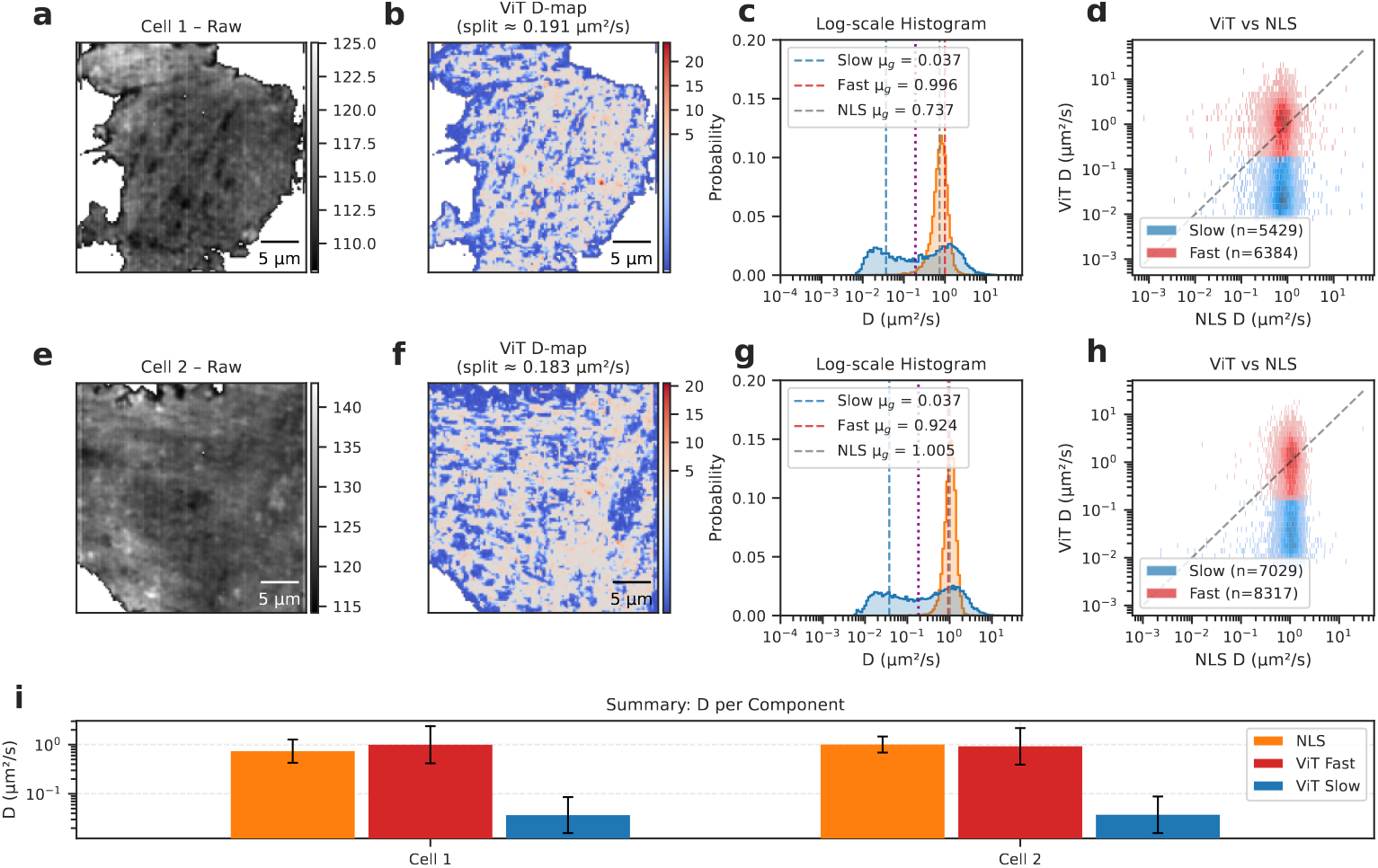
(**a, e**) Background-masked raw images for Cell 1 and Cell 2. (**b, f**) ViT *D* maps with a diverging colour scale centred on the Otsu split point, separating fast (red) and slow (blue) diffusing regions. (**c, g**) Log-scale histograms of per-pixel *D* values for ViT (blue) and NLS (orange). Vertical lines mark the geometric means of the fast (red) and slow (blue) ViT components, the Otsu split point (purple), and the NLS geometric mean (grey). (**d, h**) Two-dimensional histograms of per-pixel ViT versus NLS *D* values, coloured by Otsu classification, with the identity line (dashed). (**i**) Summary barplot of geometric mean *D* for each component across both cells. All bars represent geometric mean *×/÷* geometric standard-deviation factor.

It is important to stress that the spatial bimodality of the ViT D estimates is not identical to NLS-based fitting with 2 components. ImFCSViT is trained on homogeneous simulations and outputs a single D per pixel. Rather, the two populations reflect spatial heterogeneity in membrane diffusion, analogous to the physically distinct fast (SLB) and slow (vesicle) populations observed in the DOPC measurements. The NLS analysis, by contrast, produces a unimodal distribution whose geometric mean tends towards the ViT-derived fast component, once again demonstrating the bias towards the high-frequency fluctuations of the faster, freely diffusing population, thus leaving it unable to capture the slower hindered states. With the spatial information granted through the 3 *×* 3 window, ImFCSViT is better equipped to capture these slower states, revealing spatial heterogeneity otherwise missed through standard FCS fitting.

Furthermore, ImFCSViT results are remarkably consistent. Despite differences in cell morphology, the Otsu split thresholds were nearly identical across both cells (0.189 and 0.179 *µm*^2^*/s*), and the fast-component geometric means differ by less than 5% (0.99 and 0.95 *µm*^2^*/s*). These values are consistent with expectations of freely-diffusing membrane-anchored proteins in the plasma membrane of mammalian cells (*∼* 0.5 − 1*µm*^2^*/s*).

Beyond yielding average values, ImFCSViT simultaneously produces spatially resolved *D* maps. The fast and slow populations are not randomly intermixed but exhibit coherent spatial organization (Figure 4b,f): fast-diffusing pixels dominate extended regions of the membrane, while slow pixels cluster in discrete patches, potentially reflecting hindered diffusion within membrane microdomains modified by the underlying cytoskeleton (44), though a definitive structural analysis is beyond the scope of the current study. ViT-based diffusion coefficient-estimates thus offer a potential tool for probing the spatial organisation of membrane diffusion in living cells. Critically, ViT-based data fitting achieves this separation using only 2,500 frames (*∼* 5 s of acquisition at 2.06 ms frame time), compared to the general standard of 50 000 frames (*∼* 100 s at 2.06 ms frame time) required by NLS, making it practical for studying dynamic cellular processes on biologically relevant timescales.

### Sub-Pixel Heterogeneity

In the previous sections, our canonical 3 *×* 3 ImFCSViT model demonstrated strong capabilities in both supported lipid bilayers and live-cell plasma membranes. However, these systems are relatively homogeneous. In reality, many intracellular environments are highly ordered at the nanoscale, exposing spatial non-homogeneity that violates the assumptions of our simulation-led training process, where all training samples have presumed spatial homogeneity across all 9 pixels within the 3 *×* 3 region of interest. To evaluate the limitations of our canonical model, we probed intracellular actin fiber dynamics using CHO-K1 cells expressing F-tractin-mApple (Appendix B.2).

Actin filaments exist at structural scales of approximately 7 nm, well below the diffraction limit of standard fluorescence microscopy. Consequently, a single camera pixel can simultaneously contain stationary/hindered on-fibre states and freely-diffusing cytosolic (off-fibre) states. To establish local structural ground-truths, we use the process described in (45), utilizing Super-Resolution Radial Fluctuations (SRRF) (46) to identify on-fibre, off-fibre, and spatially mixed regions (Figure 5e).

**Figure 5:**
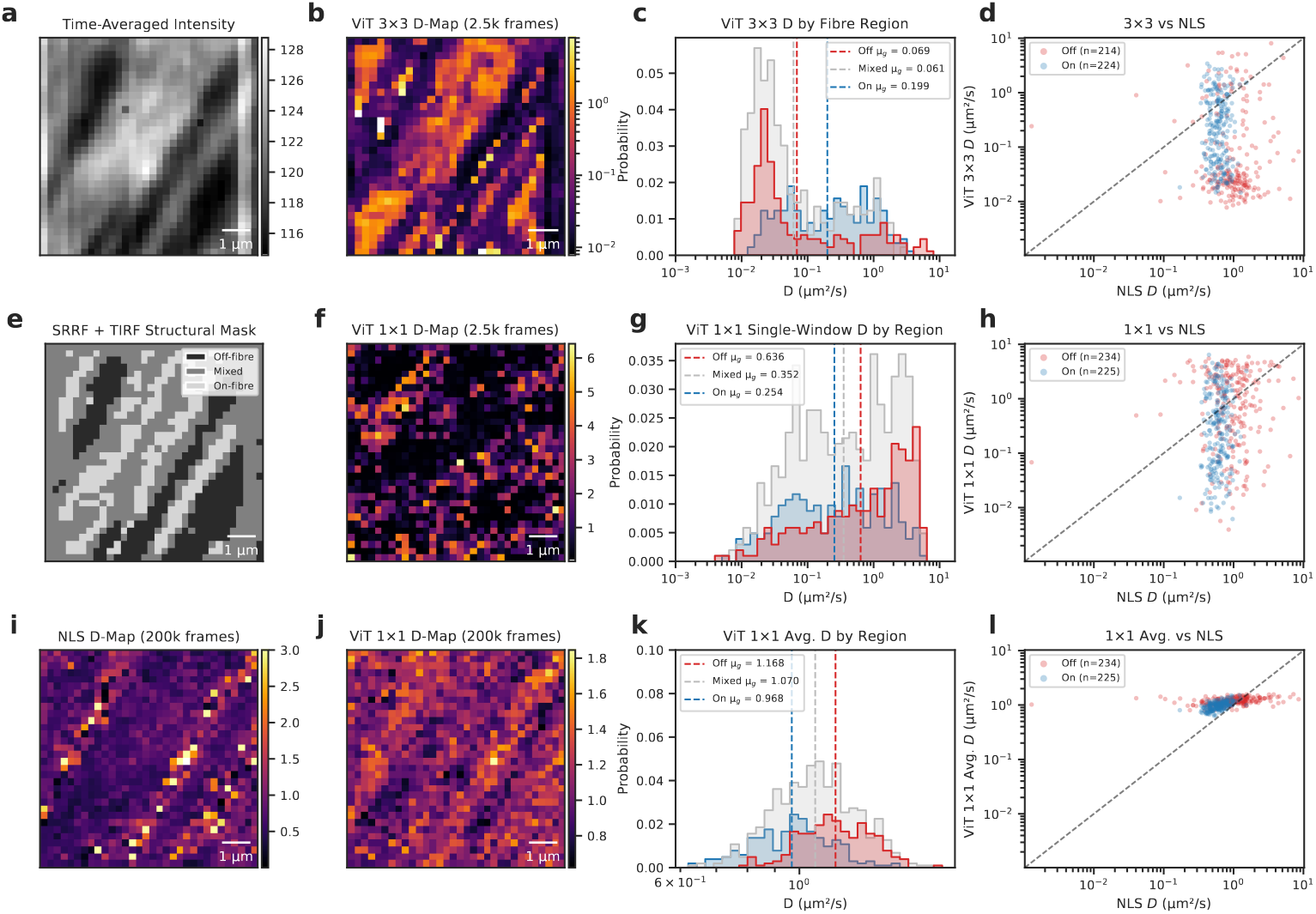
Evaluating model dependence on spatial homogeneity assumptions using F-tractin expressing cells. (**a**) Time-averaged raw intensity image. (**b**) Spatial *D*-map produced by the canonical 3×3 ViT model, displayed on a logarithmic colour scale to accommodate extreme outlier values. (**c**) Probability density of the 3 *×* 3 ViT *D* values separated by SRRF/TIRF structural mask classifications into off-fibre (red), mixed (grey), and on-fibre (blue) regions. Dashed vertical lines indicate the geometric means. (**d**) Per-pixel scatter plot of 3 *×* 3 ViT *D* versus NLS *D* for off-fibre and on-fibre classified pixels, with the identity (*y* = *x*) line. (**e**) Structural mask derived from SRRF and TIRF imaging, classifying each pixel as off-fibre, mixed, or on-fibre (from (45)). (**f**) Single-window *D*-map from the 1 *×* 1 ViT model (linear scale). The map is dominated by shot noise, obscuring spatial structure. (**g**) Probability density of 1 *×* 1 *D* values by structural region, as in (c). (**h**) Per-pixel scatter plot of 1 *×* 1 ViT *D* versus NLS *D*, as in (d). (**i**) NLS *D*-map for reference (clipped to 3*µm*^2^/*s* for visualization). (**j**) Temporally averaged single-pixel ViT *D*-map, computed as the geometric mean over 80 independent 2,500-frame windows across the full set of 200 000 frames. (**k**) Probability density of temporally averaged single-pixel ViT *D* values by region, showing the expected trend (off-fibre > mixed > on-fibre) with greatly reduced variance compared to single-window estimates. (**l**) Per-pixel scatter of temporally averaged single-pixel ViT *D* versus NLS *D*. All ViT predictions use a 2,500-frame (*∼* 5.2 s at 2.06 ms frame time) window; NLS fits use a 200 000-frame (*∼* 412 s) window.

Biologically, we expect molecules constrained by the actin filaments (on-fibre) to exhibit slower diffusion dynamics compared to the freely diffusing cytosolic regions (off-fibre). However, as shown in Figure 5c and 5d, the canonical 3 *×* 3 ImFCSViT fails to resolve this expectation. When mapping the 3 *×* 3 predictions against the SRRF-derived structural mask, the network is unable to separate the compartments and inverts the expected ordering, predicting lower mobility off-fibre (*µ*_*g*_= 0.068 *µm*^2^/*s*) than on-fibre (*µ*_*g*_ = 0.199 *µm*^2^/*s*), though with extremely large spread.

This failure is a consequence of the spatial priors embedded into the network through our simulations. When evaluating 3 *×* 3 regions that include sub-pixel resolution structures and mixtures of different diffusive components, these represent out-of-distribution (OOD) cases that the model has not encountered during training, thus resulting in predictions that are unreliable, highlighting a fundamental fragility of the canonical 3 *×* 3 ImFCSViT model.

There are two possible routes to reconciling these differences. The first is to increase the complexity of our simulations, incorporating spatial non-homogeneity that better reflects the expected data distribution of true experimental samples. However, this requires a more thorough understanding of underlying physics and biology, and also presents data labeling ambiguity (in the case where a sample contains a combination of a slow and fast diffusing region, it is ambiguous as to whether the assigned *D* label should correspond to the fast or slow region, or an average of the two). Alternatively, we can relax the spatial constraints of the ViT model by reducing the input field of view (FoV) to a single pixel.

Historically, adapting deep learning models to different input dimensions has been non-trivial. With convolutional architectures like ImFCSNet (7), reducing the input size requires a complete redesign of the network architecture and its pooling layers. In contrast, the ViT architecture processes data as a sequence of discrete tokens, making the change straightforward. By simply adjusting our tokenization strategy to tokenize 1 *×* 1 tubelets and adjusting the model embedding size, we relax the spatial constraints without altering the core transformer topology. More details are provided in Appendix D.1.

Evaluating the 1 *×* 1 ImFCSViT variant on a 2500-frame window of the same F-tractin dataset shows the recovery of the expected dynamics, identifying the fastest diffusion in off-fibre regions (*µ*_*g*_= 0.636 *µm*^2^/*s*) and the slowest dynamics localized on the actin filaments (*µ*_*g*_= 0.254 *µm*^2^/*s*) (Figure 5g). However, the resulting spatial *D*-map (Figure 5f) is visibly noisy and fails to show structural variation as seen in the NLS maps from 200 000 frames (Figure 5i). This is expected, as the intensity traces from a single pixel across 2500 frames means the input signal is dominated by shot noise. Unlike the 3 *×* 3 ImFCSViT, the 1 *×* 1 variant lacks the additional spatial information granted through the larger FoV, effectively operating on 9*×* less data than the 3 *×* 3 ViT, leading to highly variable, long-tailed distributions (*σ*_*g*_ > 5). However, as evidenced by the separation of the probability distributions in Figure 5g, the single-pixel 1 *×* 1 ViT successfully isolates the distinct dynamics of the off, mixed, and on-fibre regions using only *∼* 5 seconds (2500 frames at 2.06 ms frame time) of data.

While the 2,500-frame window successfully extracts the compartmentalized *D* distributions, generating a high-fidelity spatial map requires suppressing the high stochastic variance inherent to single-pixel, short-window traces. To demonstrate that these highly variable predictions are physically consistent and converge to the expected steady-state dynamics, we evaluate the 1 *×* 1 ImFCSViT sequentially across the full set of 200,000 frames (corresponding to *∼* 412 seconds), generating a stack of 80 localized parameter maps. By then calculating the pixel-wise geometric means across time, we integrate out the short-term fluctuations and shot noise, and obtain parameter maps that properly reflect the underlying biology, visually tracking the F-tractin filaments (Figure 5j) and producing a clean, structured image comparable to the spatial clarity of the corresponding 200 000-frame NLS map (Figure 5i).

Statistically, time-integration collapses the wide per-window variance dramatically, narrowing the per-region probability distributions (Figure 5k). The resulting data confirms the stepwise drop in molecular mobility corresponding to spatial confinement: descending from off-fibre cytosolic regions (*µ*_*g*_ = 1.167*µm*^2^/*s*), to mixed pixels (*µ*_*g*_ = 1.070*µm*^2^/*s*), and ending at on-fibre structures (*µ*_*g*_ = 0.967*µm*^2^/*s*). Although the absolute values are slightly higher than the NLS estimates (possibly due to the idealized noise conditions used during training), the rank order and spatial pattern are in good agreement.

In this case study, we demonstrate two complementary variations of the ImFCSViT framework. First, the canonical 3 *×* 3 ImFCSViT model’s ability to incorporate spatial information via its 9-pixel FoV is a powerful prior that provides a distinct advantage over standard Imaging FCS in low-frame count regimes. However, this also enforces an assumption of spatial homogeneity, which becomes a liability in highly structured environments. As demonstrated in the F-tractin-mApple case with sub-pixel heterogeneity, the 3 *×* 3 ImFCSViT produces unreliable parameter estimates. To solve this issue, we propose a single-pixel 1 *×* 1 ImFCSViT, a straightforward modification to the transformer model that relaxes the spatial constraints without requiring any further architectural changes.

The single-pixel ViT, once temporally averaged, approaches the spatial clarity of conventional NLS and faithfully reproduces the expected compartment-wise mobility trend, albeit with a positive systematic bias. We further examine the behaviour of the single-pixel ViT on homogeneous systems (DOPC supported lipid bilayers) in Appendix D.1, showing that in those cases the canonical 3 *×* 3 model remains superior, highlighting the trade-off between spatial averaging and robustness to heterogeneity.

### Runtime

Beyond its predictive capabilities, transitioning from a bespoke convolutional architecture to a standardized ViT fundamentally alters the computational profile of the inference pipeline. To quantify how theoretical efficiency translates to real-world throughput, we benchmarked the inference speeds of both our ImFCSViT variants against the ImFCSNet baseline (7). Benchmarks were run on a standard consumer-grade laptop (AMD Ryzen 7 5800H CPU, NVIDIA GeForce RTX 3060 Mobile GPU with 6GB VRAM), and all models evaluate a single 128 *×* 128 pixel image stack of 2500 frames. For ImFCSNet and the 3 *×* 3 ImFCSViT, the image is processed patch-by-patch with a 3×3 sliding window of stride 1. Meanwhile, the 1 *×* 1 ImFCSViT evaluates every pixel independently.

Crucially, it should be noted that both ImFCSViT variants provide 4 biophysical parameters in a single forward pass, whereas ImFCSNet only predicts *D*. While the 3 *×* 3 ImFCSViT model appears lighter than ImFCSNet, ViT model runtimes are generally slower than ImFCSNet. Under optimal throughput configurations (Batch Size = 128), ImFCSNet finishes slightly faster than the 3 *×* 3 ViT (4.91 s vs 5.36 s), despite ImFCSViT requiring 60% fewer operations (119.6 vs 307.3 MFLOPs). This discrepancy arises because Vision Transformers are heavily memory-bandwidth bound due to multi-head attention and LayerNorm operations, whereas spatial convolutions (ImFCSNet) utilize inherently faster cuDNN memory caching. Moreover, the fact that both models take roughly 5 seconds reflects that end-to-end inference for large datasets in this domain is ultimately bottlenecked by system data-loading (I/O) overhead and CPU-side memory allocation rather than pure GPU mathematical limits. Notably, on CPUs, the absolute runtime converges to below one minute, making deep learning-based analysis practical on standard laptop hardware, even in CPU-only mode.

Similarly, the 1 *×* 1 ImFCSViT model is dramatically lighter at 3.9 MFLOPs, reflecting the proportional decrease in the model’s embedding dimension (from 96 in the 3 *×* 3 ViT down to 24 in the 1 *×* 1 ViT). Because the 1 *×* 1 ViT tokenizes the input in 1 *×* 1 *×* 20 chunks (20 pixels per token) compared to the 3 *×* 3 *×* 10 chunks (90 pixels) of the canonical model, the required internal feature representation shrinks accordingly, speeding up computation due to the quadratic scaling of self-attention. A full discussion of tokenization scaling is provided in Appendix D.1. The 1 *×* 1 model requires only ≈1.41 s to process the 128 *×* 128 grid for a 2500-frame sequence with a GPU, and just ≈8.05 s on a CPU. This translates to an 3.48-fold speedup over ImFCSNet’s GPU time for a single map.

Crucially, the 2500-frame operating window of our networks allows the ImFCSViT models to resolve time-dependent dynamics that conventional FCS, with its 50 000-frame averaging, cannot resolve. For a typical 50 000-frame acquisition, both ImFCSViT networks can generate 20 separate parameter maps by partitioning the acquired image stack into sequential, non-overlapping windows along the time axis, producing a 20-point kinetic trace per region in approximately 20 *×* 5.3 ≈ 100 s (≈ 2 min) on the test GPU.

## DISCUSSION

In this work, we propose ImFCSViT, adopting the standard Vision Transformer (ViT) as a multi-output framework capable of simultaneously extracting multiple biophysical parameters (*D, N, B, σ*_0_) directly from raw, short-horizon fluorescence image stacks. Using only 2500 frames (2.5 s with a 1 ms frame time), a 20-fold reduction compared to the 50 000-frame acquisition typically required for conventional FCS, the ImFCSViT framework yields diffusion coefficients whose fast-diffusing component matches conventional FCS estimates while additionally estimating *N, B*, and *σ*_0_. The spatial context of the 3 *×* 3 input patch further allows ImFCSViT to uncover distinct sub-populations in both lipid bilayers and live-cell plasma membranes.

These localized sub-populations represent a crucial biologically-relevant capability, demonstrating ImFCSViT’s ability to probe spatially heterogeneous phenomena that are either invisible to, or averaged out by long-time-scale correlation functions in classical FCS. Many crucial dynamic cellular processes occur on timescales much faster than the 50 seconds required by conventional fitting. By compressing the observation window to 2.5 seconds and utilizing the raw image stacks, as opposed to the lossy ACF representation, the ImFCSViT approach enables the mapping of slow-diffusing patches in live cells. This approach provides a new method of observing dynamic spatiotemporal non-homogeneity, allowing biologists to not only gather insights that are inaccessible to standard bulk-averaged FCS, but potentially observe transient, short-lived diffusive modes.

A major practical benefit of adopting Vision Transformers is the architectural flexibility. Unlike rigid convolutional networks, which require complete architectural re-engineering to change spatial pooling, ViTs cleanly adapt to single-pixel inputs by simply altering sequence tokenization. Notably, the two configurations offer complementary properties: the 3 *×* 3 variant exploits spatial aggregation to suppress shot noise and can even decouple *D* from *σ*_0_ (Appendix A.1), while the 1 *×* 1 variant resolves sub-pixel spatial heterogeneity at the cost of higher stochastic variance, which can be overcome by temporal averaging. Crucially, one can utilize both the spatiotemporal 3 *×* 3 ImFCSViT and the 1 *×* 1 ImFCSViT configurations in tandem, making use of the strengths of each model simultaneously.

We further leverage the flexibility by training models on shorter input lengths in Appendix D.2, where we find that it is possible to further reduce the required measurement times down to *∼* 1 second (1000 frames at a 1.06 ms frame time) while still managing to accurately determine the diffusion coefficient despite a loss in precision. Even at extremes of *∼* 0.1 seconds (100 frames), ImFCSViT-based data fitting, while failing to accurately quantify exact diffusion coefficients, manages to identify large-scale shifts, thus showing potential for screening applications where change detection, rather than precise quantification, is needed. Overall, this points to our simulation-led training framework as a potential integration pathway, allowing for measurement strategies that bring us closer to true simultaneous determination of structure and millisecond molecular dynamics proposed by (32), leveraging deep learning to sidestep the minute-long measurement times required by conventional FCS-based dynamics estimation.

Beyond architectural flexibility, the ViT architecture’s self-attention mechanism also allows us to probe the network’s internal representations (Appendix E). Using simulations, we examine the self-attention weights across the final transformer block, and we observe that one head shows inter-token attention decay profiles that mirror the shape of classical ACFs, even shifting predictably in response to varying diffusion coefficients. Crucially, this emergent behaviour is not forced, as the network never has access to the ACF during training, suggesting that the sequence-learning mechanism naturally discovers methods that exploit temporal correlations through a purely data-driven representation learning process.

Despite these gains, several limitations of the present approach point to future developments. First, although ImFCSVIT reliably captures the qualitative scaling of particle number *N* and brightness *B* with concentration and laser power in DOPC bilayers, the absolute brightness estimates are not yet accurate enough for stoichiometric analysis. As detailed in Appendix A.2, when evaluated on known dimer-to-monomer populations, the network fails to recover the exact empirical brightness ratios expected by standard N&B analysis, predicting a ratio of *∼* 1.3 instead of the expected *∼* 2. While our normalization token successfully preserves the relative bulk statistics (mean and variance) necessary to track qualitative concentration changes, extracting absolute quantitative stoichiometry requires rigorous accounting of background offset and read-noise variance, which our current architecture lacks. Thus, a possible solution is noise-aware training, where camera-specific noise models and auxiliary loss components that mimic the corrections used in conventional N&B are incorporated into the training process.

Second, we choose simulation-driven training as it is the only viable path to obtaining a sufficiently large dataset with known ground truth labels. However, this does not divorce our models from approximations; rather, it simply moves the assumptions from the analytical fitting procedures up to the data generation process. Consequently, the model inherits the blind spots of the simulation design, and any phenomena not included cannot be captured. As demonstrated with the F-tractin measurements, when spatial homogeneity priors are broken through the existence of sub-pixel actin filaments, the simulation-trained 3 *×* 3 ImFCSViT fails to generalize across the simulation-to-experiment gap.

Increasing the complexity of our simulations could be a path to more generalizable models, but this requires a complete understanding of the underlying physical system, which might not be accessible. An alternative is to train on experimental data labelled by conventional fitting. However, this shifts the task from representational learning, where it organizes a high-dimensional feature representation optimized strictly for the extraction of the target properties, to functional approximation, where the network inherits the limitations inherent to the classical estimator and the lossy ACF representation.

A natural, non-circular solution to this bottleneck is self-supervised learning (SSL), where training learns representations whilst forgoing labels entirely (47). Future approaches can utilize SSL methods to train on massive datasets of unlabelled, heterogeneous experimental acquisitions, learning representations that account for noise characteristics, sample heterogeneity, and other complexities of real-world biological fluorescence. By learning these underlying distributions directly from experimental data, SSL creates robust feature representations that natively account for complex intracellular backgrounds and noise profiles. A subsequent fine-tuning step with a targeted set of simulated data would then map these robust features onto physical parameters, allowing the network to bypass the noise-related pitfalls of a purely simulation-based approach. By reducing the reliance on simulations, SSL is a viable pathway to solving the train-to-test-time distribution shifts. Notably, many modern SSL methods are transformer-specific, including the reconstruction-based MAE (35, 48) and contrastive DINO (49, 50), a further benefit to the adoption of the ViT architecture.

Looking ahead, the ImFCSViT framework can be progressively improved by richer simulations that include multi-component dynamics and binding kinetics, by noise-aware training that reconciles the competing demands of robust diffusion and accurate brightness estimation, and by SSL pre-training that grounds the model in real data distributions. Perhaps most importantly, the use of a standard, well-supported architecture means that each of these extensions can be pursued without reinventing the core network, ensuring that progress in deep learning translates directly into progress in fluorescence fluctuation spectroscopy.

## AUTHOR CONTRIBUTIONS

T.W. and S.R.S. designed the research. S.R.S. developed the ViT-based deep learning framework and conducted data analysis. Z.W., S.P. and S.Z. performed experiments. All authors contributed to the final version of the manuscript.

## ACKNOWLEDGMENTS

T.W. gratefully acknowledges funding from the Singapore Ministry of Education (T2EP30223-0042) and the Institute for Digital Molecular Analytics and Science (C-33-18-279-V12). S.R.S., Z.W., S.P., and S.Z. are recipients of NUS research scholarships.

## DECLARATION OF GENERATIVE AI AND AI-ASSISTED TECHNOLOGIES IN THE WRITING PROCESS

During the preparation of this work the author(s) used Deepseek and Gemini language models via official APIs in order to assist with proofreading and tone refinement to better reflect the expectations of academic writing. After using this tool/service, the author(s) reviewed and edited the content as needed and take(s) full responsibility for the content of the publication.

## SUPPLEMENTARY MATERIAL

An online supplement to this article can be found by visiting BJ Online at http://www.biophysj.org.

## REFERENCES

1. Douglas Magde, Elliot L. Elson, and Watt W. Webb. “Fluorescence Correlation Spectroscopy. II. An Experimental Realization”. In: Biopolymers 13.1 (1974), pp. 29±61. ISSN: 1097-0282. DOI: 10.1002/bip.1974.360130103.

2. Alexei Tcherniak et al. “Fluorescence Correlation Spectroscopy: Criteria for Analysis in Complex Systems”. In: Analytical Chemistry 81.2 (2009), pp. 746±754. ISSN: 0003-2700. DOI: 10.1021/ac8013109.

3. Jagadish Sankaran et al. “Accuracy and Precision in Camera-Based Fluorescence Correlation Spectroscopy Measurements”. In: Analytical Chemistry 85.8 (2013), pp. 3948±3954. ISSN: 0003-2700, 1520-6882. DOI: 10.1021/ac303485t.

4. Nathan Quiblier et al. “Enhancing Fluorescence Correlation Spectroscopy with Machine Learning to Infer Anomalous Molecular Motion”. In: Biophysical Journal 124.5 (2025), pp. 844±856. ISSN: 0006-3495. DOI: 10.1016/j.bpj.2025.01.026.

5. Alva Seltmann et al. “Neural Network Informed Photon Filtering Reduces Fluorescence Correlation Spectroscopy Artifacts”. In: Biophysical Journal 123.6 (2024), pp. 745±755. ISSN: 0006-3495, 1542-0086. DOI: 10.1016/j.bpj.2024.02.012.

6. Kun Zang, Zezhong Wang, and Meilisha Xu. “From Neural Networks to Transformers: Achieving High and Fast Precision in Fluorescence Correlation Spectroscopy Analysis”. In: ACS Omega 10.23 (2025), pp. 25091±25101. DOI: 10.1021/acsomega.5c03605.

7. Wai Hoh Tang et al. “Deep Learning Reduces Data Requirements and Allows Real-Time Measurements in Imaging FCS”. In: Biophysical Journal 123.6 (2024), pp. 655±666. ISSN: 0006-3495. DOI: 10.1016/j.bpj.2023.11.3403.

8. Thorsten Wohland et al. “FCS Videos: Fluorescence Correlation Spectroscopy in Space and Time”. In: Biochimica et Biophysica Acta (BBA) - General Subjects 1868.11 (2024), p. 130716. ISSN: 0304-4165. DOI: 10.1016/j.bbagen.2024.130716.

9. Michelle A. Digman et al. “Mapping the Number of Molecules and Brightness in the Laser Scanning Microscope”. In: Biophysical Journal 94.6 (2008), pp. 2320±2332. ISSN: 0006-3495. DOI: 10.1529/biophysj.107.114645.

10. Alexey Dosovitskiy et al. An Image Is Worth 16x16 Words: Transformers for Image Recognition at Scale. 2021. DOI: 10.48550/arXiv.2010.11929. arXiv: 2010.11929 (cs).

11. Douglas Magde, Elliot Elson, and Watt W Webb. “Thermodynamic Fluctuations in a Reacting SystemÐMeasurement by Fluorescence Correlation Spectroscopy”. In: Physical review letters 29.11 (1972), p. 705.

12. Rudolf Rigler and Elliot S. Elson. Fluorescence Correlation Spectroscopy: Theory and Applications. Ed. by F. P. Schäfer, J. P. Toennies, and W. Zinth. Vol. 65. Springer Series in Chemical Physics. Berlin, Heidelberg: Springer, 2001. ISBN: 978-3-642-64018-6 978-3-642-59542-4. DOI: 10.1007/978-3-642-59542-4.

13. David M. Jameson, Justin A. Ross, and Joseph P. Albanesi. “Fluorescence Fluctuation Spectroscopy: Ushering in a New Age of Enlightenment for Cellular Dynamics”. In: Biophysical Reviews 1.3 (2009), pp. 105±118. ISSN: 1867-2450. DOI: 10.1007/s12551-009-0013-8.

14. Ryan A. Colyer et al. “High-Throughput FCS Using an LCOS Spatial Light Modulator and an 8 × 1 SPAD Array”. In: Biomedical Optics Express 1.5 (2010), pp. 1408±1431. ISSN: 2156-7085. DOI: 10.1364/BOE.1.001408.

15. Ryan A. Colyer et al. “Ultra High-Throughput Single Molecule Spectroscopy with a 1024 Pixel SPAD”. In: Proceedings of SPIE 7905 (2011), pp. 10.1117/12.874238. ISSN: 0277-786X. DOI: 10.1117/12.874238.

16. Meike Kloster-Landsberg et al. “Cellular Response to Heat Shock Studied by Multiconfocal Fluorescence Correlation Spectroscopy”. In: Biophysical Journal 103.6 (2012), pp. 1110±1119. ISSN: 0006-3495, 1542-0086. DOI: 10.1016/j.bpj.2012.07.041.

17. Hans Blom et al. “Parallel Fluorescence Detection of Single Biomolecules in Microarrays by a Diffractive-Optical-Designed 2 × 2 Fan-out Element”. In: Applied Optics 41.16 (2002), pp. 3336±3342. ISSN: 2155-3165. DOI: 10.1364/AO.41.003336.

18. Balakrishnan Kannan et al. “Electron Multiplying Charge-Coupled Device Camera Based Fluorescence Correlation Spectroscopy”. In: Analytical Chemistry 78.10 (2006), pp. 3444±3451. ISSN: 0003-2700. DOI: 10.1021/ac0600959.

19. Thorsten Wohland, Sudipta Maiti, and Radek MachánÏ. An Introduction to Fluorescence Correlation Spectroscopy. IOP Publishing, 2020. ISBN: 978-0-7503-2080-1.

20. Nancy L. Thompson. “Fluorescence Correlation Spectroscopy”. In: Topics in Fluorescence Spectroscopy: Techniques. Ed. by Joseph R. Lakowicz. Topics in Fluorescence Spectroscopy. Boston, MA: Springer US, 1999, pp. 337±378. ISBN: 978–0-306-47057-8. DOI: 10.1007/0-306-47057-8_6.

21. Donald W. Marquardt. “An Algorithm for Least-Squares Estimation of Nonlinear Parameters”. In: Journal of the Society for Industrial and Applied Mathematics 11.2 (1963), pp. 431±441. ISSN: 0368-4245, 2168-3484. DOI: 10.1137/0111030.

22. Nirmalya Bag et al. “Calibration and Limits of Camera-Based Fluorescence Correlation Spectroscopy: A Supported Lipid Bilayer Study”. In: Chemphyschem : a European journal of chemical physics and physical chemistry 13.11 (2012), pp. 2784±2794. ISSN: 1439-7641. DOI: 10.1002/cphc.201200032.

23. Jonas Ries, Eugene P. Petrov, and Petra Schwille. “Total Internal Reflection Fluorescence Correlation Spectroscopy: Effects of Lateral Diffusion and Surface-Generated Fluorescence”. In: Biophysical Journal 95.1 (2008), pp. 390±399. ISSN: 0006-3495, 1542-0086. DOI: 10.1529/biophysj.107.126193.

24. Jay R. Unruh and Enrico Gratton. “Analysis of Molecular Concentration and Brightness from Fluorescence Fluctuation Data with an Electron Multiplied CCD Camera”. In: Biophysical Journal 95.11 (2008), pp. 5385±5398. ISSN: 0006-3495. DOI: 10.1529/biophysj.108.130310.

25. Anjali Gupta et al. “Heptanol-Mediated Phase Separation Determines Phase Preference of Molecules in Live Cell Membranes”. In: Journal of Lipid Research 63.6 (2022). ISSN: 0022-2275, 1539-7262. DOI: 10.1016/j.jlr.2022.100220.

26. John Kohler, Kwang-Ho Hur, and Joachim Dieter Mueller. “Autocorrelation Function of Finite-Length Data in Fluorescence Correlation Spectroscopy”. In: Biophysical Journal 122.1 (2023), pp. 241±253. ISSN: 0006-3495. DOI: 10.1016/j.bpj.2022.10.027.

27. Ryosuke Fukushima, Johtaro Yamamoto, and Masataka Kinjo. “Number and Brightness Analysis: Visualization of Protein Oligomeric State in Living Cells”. In: Advanced Imaging and Bio Techniques for Convergence Science. Ed. by Jun Ki Kim, Jeong Kon Kim, and Chan-Gi Pack. Singapore: Springer, 2021, pp. 31±58. ISBN: 978–981-336-064-8. DOI: 10.1007/978-981-33-6064-8_2.

28. H. Qian and E. L. Elson. “On the Analysis of High Order Moments of Fluorescence Fluctuations”. In: Biophysical Journal 57.2 (1990), pp. 375±380. ISSN: 0006-3495. DOI: 10.1016/S0006-3495(90)82539-X.

29. H Qian and E L Elson. “Distribution of Molecular Aggregation by Analysis of Fluctuation Moments.” In: Proceedings of the National Academy of Sciences of the United States of America 87.14 (1990), pp. 5479±5483. ISSN: 0027-8424. DOI: 10.1073/pnas.87.14.5479.

30. Joachim D Müller. “Cumulant Analysis in Fluorescence Fluctuation Spectroscopy”. In: Biophysical Journal 86.6 (2004), pp. 3981±3992. DOI: 10.1529/biophysj.103.037887.

31. Harikrushnan Balasubramanian et al. “The Dependence of EGFR Oligomerization on Environment and Structure: A Camera-Based N&B Study”. In: Biophysical Journal 121.23 (2022), pp. 4452±4466. ISSN: 0006-3495. DOI: 10.1016/j.bpj.2022.11.003.

32. Jagadish Sankaran et al. “Simultaneous Spatiotemporal Super-Resolution and Multi-Parametric Fluorescence Microscopy”. In: Nature Communications 12.1 (2021), p. 1748. ISSN: 2041-1723. DOI: 10.1038/s41467-021-22002-9.

33. Ashish Vaswani et al. “Attention Is All You Need”. In: Advances in Neural Information Processing Systems. Vol. 30. Curran Associates, Inc., 2017.

34. Salman Khan et al. “Transformers in Vision: A Survey”. In: Acm Computing Surveys 54.10s (2022), 200:1±200:41. ISSN: 0360-0300. DOI: 10.1145/3505244.

35. Zhan Tong et al. VideoMAE: Masked Autoencoders Are Data-Efficient Learners for Self-Supervised Video Pre-Training. 2022. DOI: 10.48550/arXiv.2203.12602. arXiv: 2203.12602 (cs).

36. Limin Wang et al. “VideoMAE V2: Scaling Video Masked Autoencoders with Dual Masking”. In: 2023 IEEE/CVF Conference on Computer Vision and Pattern Recognition (CVPR). 2023, pp. 14549±14560. DOI: 10.1109/CVPR52729.2023.01398.

37. Thaddäus Wiedemer et al. Video Models Are Zero-Shot Learners and Reasoners. 2025. DOI: 10.48550/arXiv.2509.20328.

38. Adrien Bardes et al. Revisiting Feature Prediction for Learning Visual Representations from Video. 2024. DOI: 10.48550/arXiv.2404.08471. arXiv: 2404.08471 (cs).

39. Georgios Tziafas and Hamidreza Kasaei. “Early or Late Fusion Matters: Efficient RGB-D Fusion in Vision Transformers for 3D Object Recognition”. In: 2023 IEEE/RSJ International Conference on Intelligent Robots and Systems (IROS). 2023, pp. 9558±9565. DOI: 10.1109/IROS55552.2023.10341422.

40. Lucas Beyer, Xiaohua Zhai, and Alexander Kolesnikov. Better Plain ViT Baselines for ImageNet-1k. 2022. DOI: 10.48550/arXiv.2205.01580. arXiv: 2205.01580 (cs).

41. Xiaokang Chen et al. “Context Autoencoder for Self-supervised Representation Learning”. In: International Journal of Computer Vision 132.1 (2024), pp. 208±223. ISSN: 1573-1405. DOI: 10.1007/s11263-023-01852-4.

42. J. L. W. V. Jensen. “Sur Les Fonctions Convexes et Les Inégualités Entre Les Valeurs Moyennes”. In: (1906). DOI: 10.1007/bf02418571.

43. Nobuyuki Otsu. “A Threshold Selection Method from Gray-Level Histograms”. In: IEEE Transactions on Systems, Man, and Cybernetics 9.1 (1979), pp. 62±66. ISSN: 2168-2909. DOI: 10.1109/TSMC.1979.4310076.

44. Shambhavi Pandey and Thorsten Wohland. “Ghosts on the Membrane: Cytoskeletal Pinning Influences Nanoscale Cell Membrane Organization”. In: Biomolecules 16.4 (2026), p. 596. ISSN: 2218-273X. DOI: 10.3390/biom16040596.

45. Shambhavi Pandey, Nithin Pathoor, and Thorsten Wohland. “Super-Resolution Algorithms for Imaging FCS Enhancement: A Comparative Study”. In: Biophysical Journal 124.20 (2025), pp. 3428±3440. ISSN: 0006-3495, 1542-0086. DOI: 10.1016/j.bpj.2025.03.031.

46. Nils Gustafsson et al. “Fast Live-Cell Conventional Fluorophore Nanoscopy with ImageJ through Super-Resolution Radial Fluctuations”. In: Nature Communications 7.1 (2016), p. 12471. ISSN: 2041-1723. DOI: 10.1038/ncomms12471.

47. Randall Balestriero et al. A Cookbook of Self-Supervised Learning. 2023. DOI: 10.48550/arXiv.2304.12210. arXiv: 2304.12210 (cs.LG).

48. Kaiming He et al. “Masked Autoencoders Are Scalable Vision Learners”. In: 2022 IEEE/CVF Conference on Computer Vision and Pattern Recognition (CVPR). 2022, pp. 15979±15988. DOI: 10.1109/CVPR52688.2022.01553.

49. Maxime Oquab et al. DINOv2: Learning Robust Visual Features without Supervision. 2024. DOI: 10.48550/arXiv.2304.07193. arXiv: 2304.07193 (cs).

50. Oriane Siméoni et al. DINOv3. 2025. DOI: 10.48550/arXiv.2508.10104. arXiv: 2508.10104 (cs.CV).

