## Supplemental Materials for "Vision Transformers as a Framework for Imaging Fluorescence Correlation Spectroscopy"

#### A ADDITIONAL RESULTS

##### A.1 Defocusing

Establishing a dynamic ground truth for the empirical point-spread-function (PSF) during live acquisition is experimentally challenging. In Imaging FCS specifically, PSF calibrations are generally done by optimizing the structure factor $\sigma_0$ such that the Imaging FCS-determined diffusion coefficient value is constant across different pixel binning amounts in a homogeneous sample (*i.e.*, the structure factor perfectly captures the PSF shape well regardless of how many pixels are convolved in the fitting process) (1). To verify ImFCSViT's $\sigma_0$ output, we use the POPC bilayer sample from (2) with an induced defocusing event to illustrate the capabilities of our model.

Here, we visualize the diffusion coefficient evolution obtained by our vision transformer against the standard FCS fitting protocol. We apply a 2500-frame sliding window for the ImFCSViT models, and a 50 000-frame sliding window for standard NLS fitting. For parity with Figure 2d from (2), we plot the arithmetic mean and standard deviation, though we stress that the training process inherently optimizes for log-space normality, which is true for the results in (2) as well. We observe that, much like (2), the $3 \times 3$ ImFCSViT is robust to defocusing, with a diffusion coefficient value that remains constant even post-defocusing, whereas the standard FCS-based fitting model interprets the increased PSF size as a slowdown in diffusion (Figure A.1a).

By visualizing the ImFCSViT model's PSF target, we observe a corresponding increase of the PSF over time from $\sim 0.78$ to $\sim 0.84$ (Figure A.1b), corresponding to the defocusing event from approximately 250 s, where deviations from the initial calibrated 0.8 value can be observed. This demonstrates that our model is able to quantify the PSF changes over time.

Crucially, the model does not rely on heavy binning routine for standard PSF calibration, meaning it unlocks a new pathway for the analysis of spatial PSF variations. However, this capability fundamentally requires the spatial context of the $3 \times 3$ input. When we evaluate the same defocusing event with the single-pixel $1 \times 1$ ImFCSViT, the model $D$ predictions drops similarly to the single-pixel NLS estimates (Figure A.1c). While we do observe the $1 \times 1$ ImFCSViT being able to estimate a slight increase of the PSF (Figure A.1d), we hypothesize that this is the parameter crosstalk between $D$ and the PSF structure factor $\sigma_0$ as with standard FCS in Equation 2, occurring because the $1 \times 1$ ImFCSViT lacks the spatial component of the $3 \times 3$ model, misrepresenting the expansion of the observation volume as a decrease in mobility.

##### A.2 Stoichiometry and N&B

To establish the ground-truth stoichiometric ratios for our mStayGold populations, standard N&B analysis was performed on the 20 ms acquisitions of 1000 frames. Following the standard procedure from (3), variance-maintaining polynomial detrending was applied with order 4 to mitigate the reduced intensity due to photobleaching. Utilizing the concurrently acquired dark image stacks, the data was corrected for intensity offsets and readout noise variance. N&B analysis was done following Equation 5 and 4. Finally, Otsu-based thresholding was done to exclude off-cell background regions from analysis. For all 10 samples (5 monomers and 5 dimers), we perform aggregation of brightness ($B$) values across each cell, obtaining the median brightness for each sample. Using standard N&B analysis, we obtain a dimer-to-monomer brightness ratio of 1.98, which matches the theoretical $2\times$ brightness ratio of mStayGold dimers.

For ImFCSViT, we approximate the N&B process by using the same training procedure discussed in Appendix C, and using the $N$ and CPS heads as a proxy for apparent particle density and molecular brightness. For the ImFCSViT estimates, we use 2500 frames acquired at 2.06 ms frame time, matching the known working acquisition settings of the CHO-K1 cells discussed in the Results section. Using the same statistical analysis, we obtain a dimer-to-monomer brightness ratio of 1.35, an

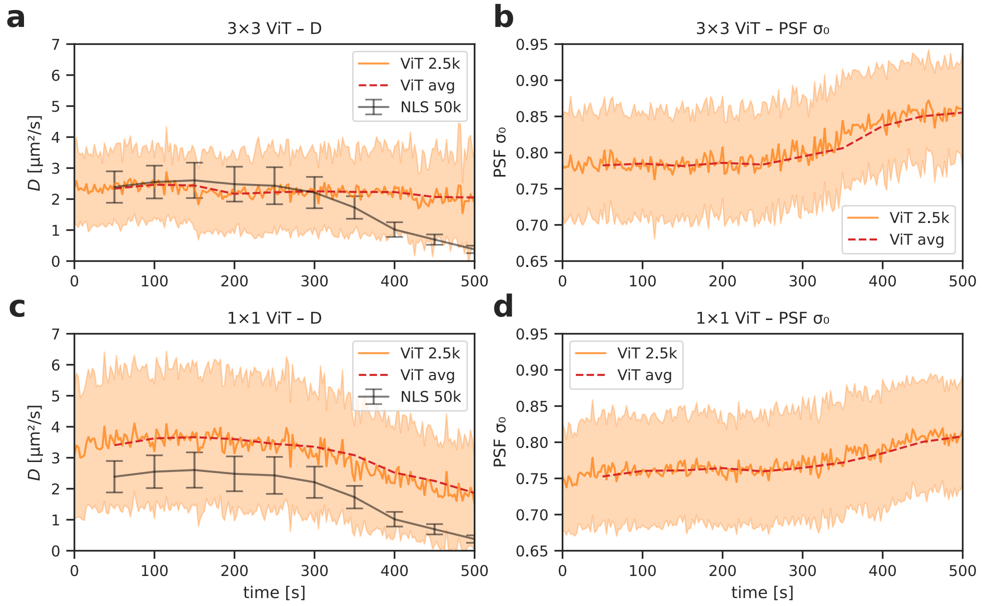


Figure A.1: Validation of ViT PSF estimation using an induced defocusing event on a POPC supported lipid bilayer. (**a**) Time-series of the diffusion coefficient $D$ extracted by the $3 \times 3$ ViT (2 500-frame sliding window, blue) and by standard NLS fitting (50 000-frame sliding window, black). The ViT estimate remains constant throughout the defocusing event, whereas the NLS estimate interprets the enlarged PSF as a slowdown in diffusion. The dashed line shows the per-file mean of the ViT predictions. (**b**) Corresponding evolution of the structure factor $\sigma_0$ from the $3 \times 3$ ViT, which increases from $\sim 0.78$ to $\sim 0.84$ as the sample is progressively defocused, matching the expected PSF broadening. (**c**) Same as (**a**) but for the $1 \times 1$ ViT. The loss of spatial context causes the model to exhibit the classic parameter crosstalk between $D$ and $\sigma_0$: the diffusion coefficient appears to decrease as the PSF grows. (**d**) PSF output of the $1 \times 1$ ViT. Although a slight upward trend is discernible, the fidelity is substantially reduced compared to the $3 \times 3$ model, highlighting the necessity of the $3 \times 3$ input patch for robust PSF characterization. In all panels, shaded bands indicate $\pm 1$ standard deviation of the ViT predictions within each 2500-frame chunk.

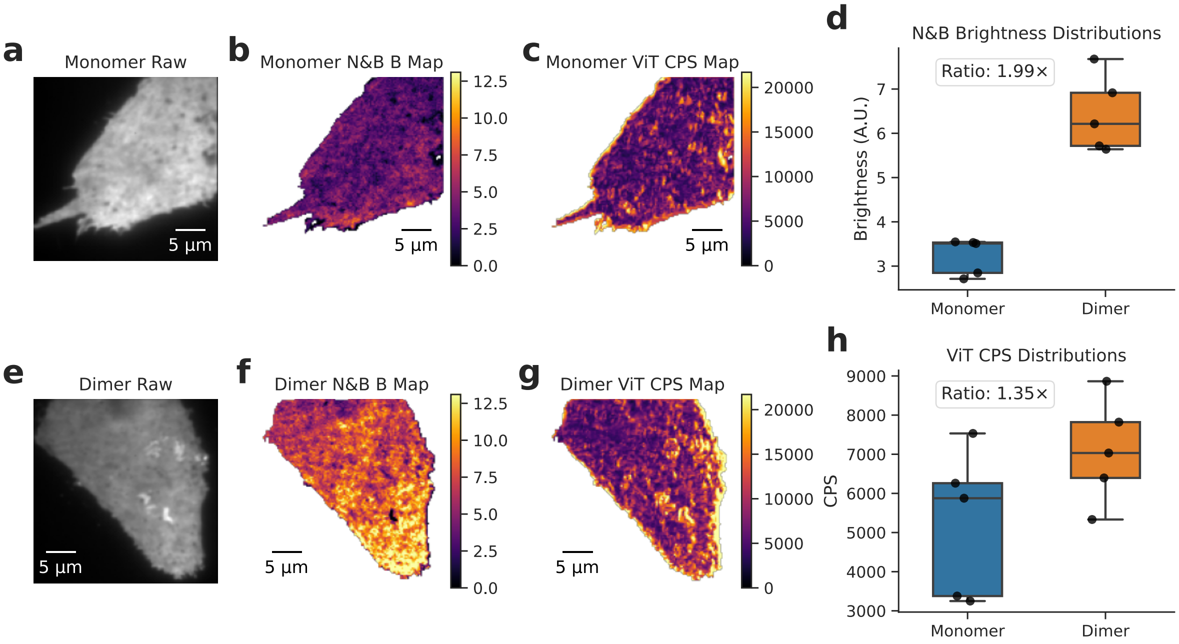


Figure A.2: Comparison of oligomeric state quantification between traditional Number and Brightness (N&B) analysis and ImFCSViT in CHO-K1 cells expressing mStayGold. (**a**-**c**) Representative field of view of a cell expressing mStayGold monomers, showing the time-averaged raw image (**a**), the N&B-derived apparent brightness map (**b**), and the ViT-inferred Counts Per Second (CPS) map (**c**). (**e**-**g**) Corresponding representative images for a cell expressing mStayGold dimers. (**d**) Distribution of median cellular N&B brightness values across the dataset for both monomeric and dimeric constructs. The annotated fold change highlights the expected relative increase in intensity per particle. (**h**) Distribution of median cellular ViT CPS values. In (**d**) and (**h**), individual points represent the spatial median value of a single image acquisition.

underestimation compared to the known true values. In Figure A.2, while ImFCSViT is able to consistently assign a higher brightness value to the dimer samples, it is unable to quantitatively extract the expected 2× ratio.

While the operating regimes of each approach are different in terms of frame time and thus signal-to-noise ratios, we emphasize that these specific regimes are established standards for each approach. To ensure that the compressed stoichiometric ratio observed in ImFCSViT predictions (1.35×) was not merely an artifact of the shorter integration time, we applied conventional N&B analysis to the same 2.06 ms acquisitions. This analysis still successfully recovered a 1.9× dimer-to-monomer brightness ratio. Thus, the observed discrepancies in stoichiometric ratios are not attributable to differences in frame time.

To investigate the failure, we fall back to simulations, simulating a set of 150 simulation samples with variable $D$ (varied in the range $[0.5, 10.0]$ in 0.5 intervals), $N$ (varied in range $[1,5]$ particles per-pixel in increments of 1) and CPS (5000, 10 000 and 15 000). We test these simulations in the noiseless case, and a case where we add simulated EMCCD noise based on the camera's dark image (noise scale = 1).

From Figure A.3a,d, we observe that the network-estimated $D$ values are consistent across both noise levels, whereas $N$ suffers in the presence of noise (Figure A.3b,e). This mirrors the sensitivity of the denominator in Equation 4 to noise, suggesting that the network, like the analytical estimator, struggles to recover clean second-moment information in low-SNR conditions. This effect is mitigated in high-concentration samples like the DOPC supported lipid bilayers, where the higher fluorescent particle density allows the effect of noise to be negated (Figure 3a-b).

In the case of the brightness predictor (CPS) (Figure A.3c, f), we observe that noise alone does not explain ImFCSViT's inability to capture the expected stoichiometric ratios. We note that these fits pool data across a range of $D$ and $N$ values; minor variations with particle density are visible in the CPS predictions but do not affect the sign or magnitude of the noise-induced intercept shift. Even in the noiseless case (Figure A.3c), ImFCSViT estimates a ratio of 1.77 between the known ground truths of 10 000 and 5000. Using linear regression, we observe a y-intercept of approximately 1431 (Figure A.3c). The origin of this offset is not fully understood; it may reflect the network's tendency to infer a minimal fluorescence level from the training distribution, or a bias inherent in the architecture's regression head, even in the presence of no noise.

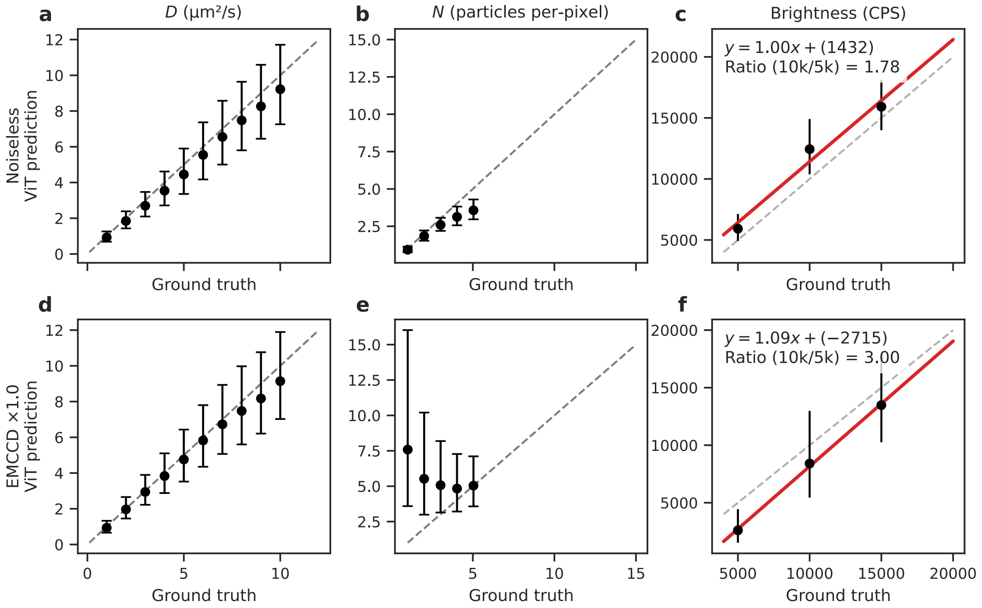


Figure A.3: ImFCSViT predictions on simulations under noiseless and EMCCD-noise conditions. (**a**) ViT-estimated diffusion coefficient $D$ versus simulation ground truth values. Geometric mean and geometric standard-deviation factor are shown for each sample. (**b**) ViT-estimated particle density $N$ versus simulation ground truth values. (**c**) ViT-estimated brightness (CPS) versus simulation ground truth values. Linear regression fit parameters and computed ratio of 10 000 CPS against 5000 CPS are shown in inset. Line of best fit shown in red. (**d-f**) Same as row 1, but with added EMCCD noise, sampled from a camera dark image as detailed in C.1.

When introducing simulated EMCCD noise, linear regression shows that the model maintains a gradient of near 1, but the intercept shifts downwards to $-2715$. While this negative intercept is an extrapolation of the linear fit (as the network's predictions are strictly positive), it illustrates that the model learns a negative bias to implicitly compensate for readout variance. Consequently, this negative penalty inflates the calculated brightness ratio under pure camera noise from 1.77 to 3.00.

However, the fact that stoichiometric ratios increase under simulated camera noise demonstrates that read-noise alone does not explain the model's in vivo failure, where brightness ratios are instead compressed. This discrepancy exposes two further limitations of our simulation design. First, in our simulations, every particle is assigned the exact same brightness value. In reality, due to stochastic fluorophore maturation, photobleaching, and transient dark states, a biological dimer population does not possess a uniform $2\times$ brightness. Instead, it exists as a mixture of species containing two, one, or zero active chromophores. Just as the model could not resolve sub-pixel diffusion heterogeneity in the F-tractin sample, it cannot resolve brightness heterogeneity. Second, our simulations lack a constant intensity background. In live-cell measurements, uncorrected cellular autofluorescence and out-of-focus signal artificially inflate the mean intensity of the trace ($\langle I \rangle$). Standard N&B explicitly corrects for this via background subtraction (Equation 5). Without this background-aware prior, the network is forced to evaluate traces containing unmodeled offsets, pushing the input outside the training distribution and further degrading the network's ability to extract the underlying variance from the signal.

Crucially, ImFCSViT in it's current form does not have access to a dark image input like standard N&B. The network's response to EMCCD noise shifts the brightness offset in a way that does not correct the ratio compression; this suggests that the model has not learned to separate shot-noise variance from true molecular brightness, a task that N&B performs explicitly via camera-noise subtraction. While the network successfully extracts dynamics in a noise-robust manner, extracting quantitative stoichiometry will require future models to incorporate noise-aware training regimens and background-corrected loss functions.

#### A.3 DOPC Bilayers

In Figure 2, we presented a single representative sample to demonstrate the feasibility of using non-parametric fast/slow-component separation to identify slow-moving residual vesicles. To confirm that this is not an isolated artifact and to demonstrate the reliability of the method, we extend this visualization across all 12 available experimental conditions (varying concentrations and illumination powers).

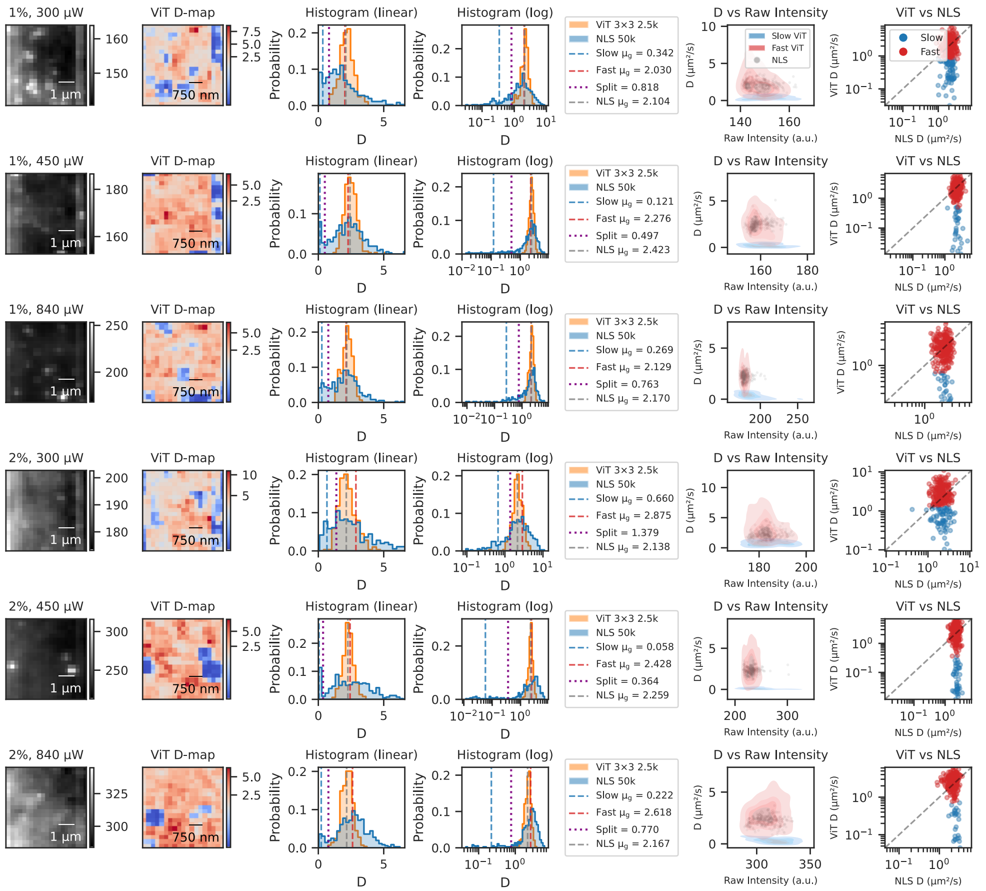


Figure A.4: Individual DOPC samples of concentrations 1% and 2% and corresponding analyses of $3 \times 3$ ImFCSViT-derived $D$ maps. From left-to-right: Raw sample visualization averaged across the first 2500 frames; The $3 \times 3$ ViT-derived $D$-map, corresponding to the first 2500 frames; Linear histogram of $D$ values with NLS (orange) and ViT (blue); Log-histogram of $D$ values with NLS (orange) and ViT (blue) and corresponding Otsu split point (purple), slow (blue) and fast (red) geometric means for the ViT, and NLS-derived geometric mean (gray); Summary statistics and corresponding values; ViT-$D$ vs. intensity map, showing the distribution of low-$D$ residual vesicle regions corresponding to higher intensity values; Scatter plot of ViT-derived $D$ values versus the NLS-fitted $D$ values, colored by fast (red) and slow (blue) pixels as determined through non-parametric Otsu.

As shown in Figures A.4 and A.5, residual vesicles are present in every measurement. The $3 \times 3$ ImFCSViT consistently isolates this slow-moving/immobile component, while simultaneously and accurately recovering the bulk membrane fast-component through the non-parametric Otsu analysis method, all while utilizing just 2500 frames.

### B MEASUREMENTS

#### B.1 DOPC and POPC Supported Lipid Bilayers

Supported lipid bilayers (SLBs) were used as model membrane systems for calibration and benchmark measurements. The primary lipid used was 1,2-dioleoyl-sn-glycero-3-phosphocholine (DOPC) (Avanti Polar Lipids, Alabaster, AL, USA). To visualize membrane dynamics, we used the head group–labeled fluorescent lipid probe 1,2-dimyristoyl-sn-glycero-3-phosphoethanolamine-N-(lissamine rhodamine B sulfonyl) (RhoPE) (Avanti Polar Lipids).

To evaluate the effect of fluorophore concentration on the deep learning parameter estimates, DOPC samples were prepared with lipid to fluorescent dye concentration ratios varying from 1% to 4%, corresponding to 50 to 200 nM of the fluorophore in 50 nM increments.

SLBs were prepared using a standard vesicle fusion protocol (4). Briefly, appropriate amounts of lipid and dye stock solutions were mixed in a clean round-bottom flask and evaporated using a rotary evaporator (Rotavap R-210, Büchi, Switzerland) for 3

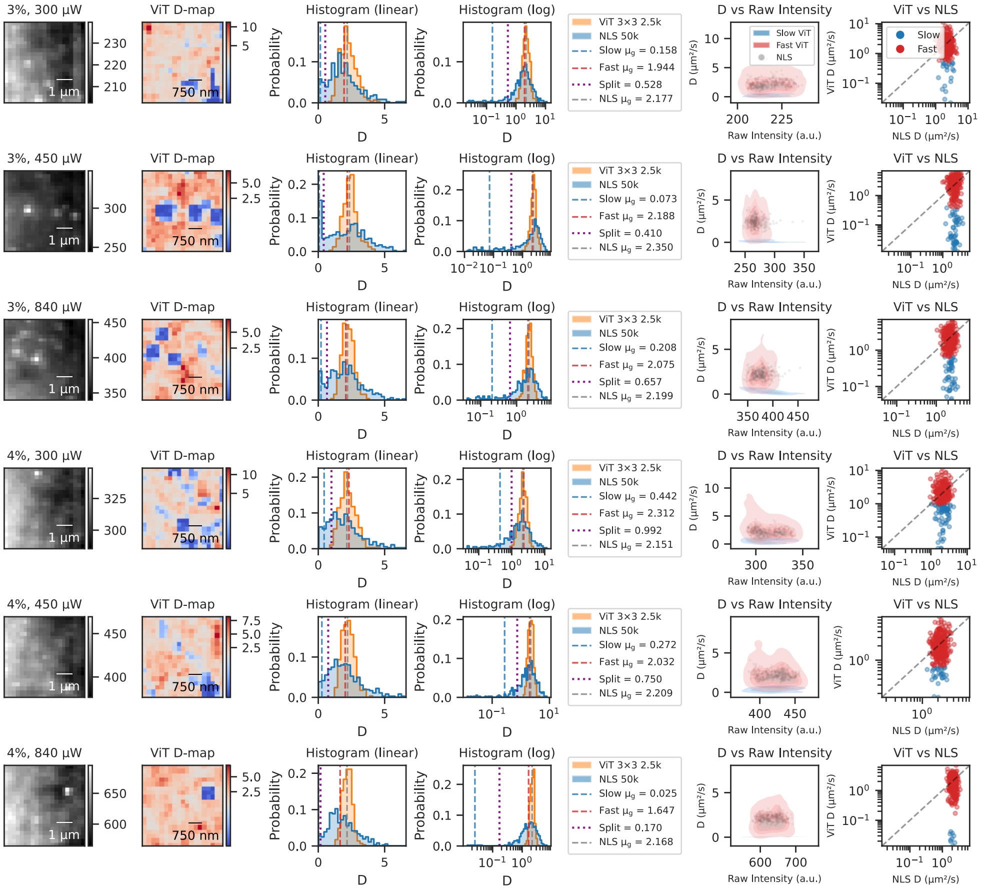


Figure A.5: Continuation of Figure A.4, with concentrations 3% and 4%.

hours to form a thin lipid film. The dried film was rehydrated in a buffer containing 10 mM HEPES and 150 mM NaCl (pH 7.4), followed by bath sonication (FB15051, Fisher Scientific, Singapore) until the solution became clear. The resulting small unilamellar vesicles (SUVs) were deposited onto cleaned glass coverslips, incubated at 65 °C for 20 min, and then cooled to room temperature for 20 min to form the bilayer. Excess unfused vesicles were removed by repeated rinsing with the same HEPES buffer.

Fluorescence fluctuations of the SLBs were recorded utilizing total internal reflection fluorescence (TIRF) microscopy. Image stacks were acquired using an electron-multiplying charge-coupled device (EMCCD) camera (iXon EM+ 860, Andor Technology, Belfast, UK) cooled to −80 °C. The camera was operated in kinetic mode with a constant baseline clamp applied to minimize baseline fluctuations, using a 10 MHz pixel readout speed and an EM gain of 300. For these lipid bilayer measurements, acquisitions were recorded at a frame time of 1.06 ms.

The POPC dataset demonstrating defocusing (Appendix A.1) was obtained from (2), with preparation and imaging details discussed accordingly.

##### B.2 CHO-K1 Cells

Live-cell measurements were performed using Chinese Hamster Ovary (CHO-K1) (CCL-61, ATCC, Manassas, VA) cells transiently expressing either a freely diffusing plasma-membrane-targeted fluorescent protein (PMT-mApple) or an actin-binding probe (F-tractin-mApple).

Briefly, CHO-K1 cells were cultured in Dulbecco's Modified Eagle Medium (DMEM, high glucose with L-glutamine, without sodium pyruvate) supplemented with 10% fetal bovine serum and 1% penicillin-streptomycin at 37 °C and 5% $CO_2$. Cells were transiently transfected at approximately 90% confluency using Lipofectamine 3000 (Invitrogen, Waltham, MA) with 100 ng of the respective plasmid constructs. Following a 36- to 48-hour incubation period, cells were washed and transferred to phenol red-free DMEM for fluorescence imaging.

Full details regarding plasmid construction, cell culture, and extended transfection protocols for the PMT-mApple and F-tractin-mApple constructs have been previously described (5, 6).

TIRF imaging was performed with a 100× oil-immersion objective (NA 1.49) at a penetration depth of 80 nm, using an illumination laser power of 1 mW. Data acquisition was performed using the same EMCCD parameters as the SLBs (−80 °C, kinetic mode, baseline clamp, 10 MHz readout, EM gain 300). For the PMT-mApple cells, stacks of 100,000 frames were recorded at a frame time of 2.06 ms.

The F-tractin-mApple dataset evaluated in this work, along with its corresponding Super-Resolution Radial Fluctuations (SRRF) structural mask used for regional classification, was obtained directly from (5) to ensure direct comparability. As described in the original study, the F-tractin dataset comprised 200,000 frames acquired at a corresponding 2.06 ms frame time.

##### B.3 mStayGold N&B Cells

To evaluate the network's ability to extract quantitative brightness ($B$) and stoichiometric ratios, live-cell control measurements were performed using CHO-K1 cells cultured as described above.

To provide known monomeric and dimeric reference populations, cells were transiently transfected to express either a monomeric plasma-membrane-targeted mStayGold (PMT-mStayGold) or its tandem dimer variant (PMT-mStayGold2). Transfection was performed via electroporation using Lipofectamine 3000 (Thermo Fisher Scientific, Waltham, MA) with 50 ng of the respective plasmid DNA. The generalized molecular cloning strategy for generating the monomer PMT-FP and dimer PMT-FP2 constructs has been detailed previously (3).

Following electroporation, cells were seeded onto glass-bottom culture dishes and incubated for 36 to 48 hours. Prior to fluorescence measurement, the cells were washed with Hanks' Balanced Salt Solution (HBSS) containing $Ca^{2+}$ and $Mg^{2+}$ and transferred to phenol red-free DMEM to minimize background autofluorescence.

To establish the baseline stoichiometric ratio via standard N&B analysis, TIRF image stacks of non-transfected cells were acquired for 1,000 frames at a 20 ms exposure time. Dark image stacks (acquired with no sample illumination) utilizing the exact spatiotemporal settings were concurrently acquired to provide the necessary offset and readout noise variance characterizations. Finally, to evaluate deep learning capabilities on the same populations, a corresponding set of 50,000-frame image stacks was acquired at a faster 2.06 ms frame time, consistent with the prior imaging parameters used in this work.

#### C TRAINING SETUP

To train our ImFCSViT models, we employed a standardized data simulation and training approach across all experiments.

#### C.1 Data Simulation

We use the simulation framework from (7) with no modifications to the underlying physical model. To speed up training, we optimized the queue system to utilize CUDA streams, allowing asynchronous simulation and training for better utilization of GPU resources. We simulate $N$ fluorescent particles randomly initialized on a 2-dimensional grid. We simulate a total of 2500 frames, each split into 30 independent simulation sub-steps. Each simulation step relocates the particle based on a 2D random walk (informed by the diffusion coefficient $D$). Photon emission is then simulated by sampling an emitted photon count from a Poisson distribution (informed by the brightness $B$ in the form of counts-per-second) distributed in a spatial Gaussian profile around the particle (spread informed by the PSF calibration factor $\sigma_0$).

Our training targets are assigned as follows:

1. $D$ as the constant global diffusion coefficient.
2. $N$ is the total number of simulated particles across the full simulation region. For the purposes of reporting, we convert the total number of particles into particle density by dividing the estimated number by the total number of pixels in the simulation area ($15 \times 15$ for our configurations).
3. Brightness $B$ is the single-particle brightness (counts-per-second per molecule) used in the simulation; all particles share the same $B$.
4. $\sigma_0$ as the global PSF structure factor term for all particles.

Simulation parameters can be split into constants, which are experiment and setup-specific (Table 3), and variables, which are user-specified and define the range of targets the model will see during training (Table 4). As discussed in the main text, we sample these variables far beyond expected biological regimes to ensure generalizability across a wide range of sample conditions. For instance, diffusion is sampled up to $D = 20$ µm$^2$/s to encompass the theoretical velocity limit observable across an expanded $3 \times 3$ convolved FoV.

Table 3: Fixed instrumental parameters used to generate the simulated training sets corresponding to the experimental datasets used.

| **Parameters** | **Lipid bilayers** | **PMT-mApple & F-tractin-mApple Cells** | **mStayGold N&B Cells** |
|---|---|---|---|
| Sensor Pixel Size ($\mu$m) | 24 | 24 | 24 |
| Objective Magnification | 100x | 100x | 100x |
| Emission Wavelength (nm) | 583 | 592 | 510 |
| PSF Calibration Factor | 0.8 | 0.96 | 0.96 |
| Numerical Aperture | 1.5 | 1.49 | 1.49 |
| Frame Time (ms) | 1.06 | 2.06 | 2.06 |

Table 4: Variable parameters used during generation of training datasets. For simplicity, all $3 \times 3$ ImFCSViT models use the exact same parameter ranges. $1 \times 1$ ViT specific modifications are discussed in Appendix D.1

| **Variables** | **Sampling Minimum** | **Sampling Maximum** | **Sampling Scale (Log/Linear)** |
|---|---|---|---|
| Diffusion Coefficient, $D$ ($\mu m^2/s$) | 0.01 | 20.00 | Log |
| Total Particles (per $15 \times 15$ grid) | 10 | 30000 | Log |
| Particle Brightness (CPS) | 500 | 20000 | Log |
| PSF Factor Spread ($\Delta\sigma_0$) | -0.2 | +0.2 | Linear |
| EMCCD Noise Scale Factor | 0.0 | 6.0 | Linear |

For all simulations, we use a grid of size $15 \times 15$, and extract the central $3 \times 3$ ROI to use for training, avoiding boundary artifacts. Our simulation harness pairs each simulation with the parameters used to generate said sample. This allows us to produce input/target pairs to be used during training. For all trained models, we expose all varied parameters as targets, excluding the EMCCD noise scale factor, which is considered to be a data augmentation strategy aiming to make our model robust to camera-specific background noise.

To model EMCCD noise, we use an empirically derived distribution based on collected dark image measurements, which ensures that our modeled EMCCD noise matches the noise profile of each experimental setup. For each experimental setup, we collect 100 000 frames with the camera shutter off and no illumination, obtaining a dark image stack that characterizes the noise profile of said experimental setup. These images are corrected as per (8) by subtracting the row-wise and column-wise means, removing systematic contributions from the EMCCD detector. The corrected dark image stack is then taken to be the characteristic noise of the system. By obtaining a histogram of pixel intensity counts across the corrected dark image stack, we obtain an empirical noise distribution that can be sampled from using inverse transform sampling.

We deliberately choose to exclude simulated Gaussian noise in our simulations, as it is not an accurate representation of the noise expected from experimental data, and empirically, is shown to result in incorrect particle density and brightness predictions.

Because parameters like $N$ and $B$ are sampled log-uniformly, the training dataset contains a high proportion of low-concentration, low-brightness samples. By pairing this low-signal bias with aggressive empirical noise scaling (up to $6\times$), we enforce the heavy regularization at the dataset-level, forcing the network to learn robust spatiotemporal feature representations rather than memorizing clean intensity fluctuations.

#### C.2 Model Design

The adoption of the Vision Transformer (ViT) stems from two main goals. First, we aimed to adhere to the canonical ViT architecture (9), ensuring that future advancements in transformer interpretability and scaling can be readily translated to deep learning in FCS. Second, we constrained the model capacity to remain below the parameter count and computational cost of the convolutional baseline (ImFCSNet (2), $\approx 0.72$M parameters). This ensures that the predictive and spatiotemporal gains demonstrated by our models are fundamentally attributable to the self-attention sequence-learning paradigm and the flexibility of the ViT architecture, rather than brute-force scaling of model capacity. To achieve this, we utilized the ViT-Tiny configuration (10) as our starting point. As detailed in Table 5, we scaled down the structural hyperparameters, specifically the embedding dimension and MLP expansion factors, formulating two discrete model variants corresponding to our input structures: the canonical $3\times3$ and single-pixel $1\times1$ ImFCSViT. The proportional scaling of the embedding dimension from 96 down to 24 for the single-pixel variant ensures optimal parameter density without structural bottlenecking (detailed further in Appendix D.1). Both variants maintain a significantly smaller footprint than standard vision models while retaining the standard multi-headed attention mechanism native to transformer models, enabling the extraction of spatiotemporal features from the raw image stacks.

Notably, the ViT architecture's ability to handle arbitrary state variables through the use of additional tokens is also crucial. Because Transformers process data as an unordered sequence, it is trivial to inject a global context vector like our normalization token alongside the spatiotemporal tokens. This is a clear benefit over rigid convolutional networks like ImFCSNet, which struggle to inject non-spatial global variables without complex architectural modifications.

Table 5: Model architecture scaling and parameter size comparison against standard baseline models.

| **Model** | **Embed. Dim.** | **Heads** | **Layers** | **MLP Exp. Factor** | **Params** | **Source** |
|---|---|---|---|---|---|---|
| $1\times1$ **ImFCSViT** | 24 | 6 | 6 | $2\times$ | 0.04M | - |
| $3\times3$ **ImFCSViT** | 96 | 6 | 6 | $2\times$ | 0.59M | - |
| ImFCSNet | - | - | - | - | 0.72M | (2) |
| ViT-Tiny | 192 | 3 | 12 | $4\times$ | 5M | (10) |
| ViT-Small | 384 | 6 | 12 | $4\times$ | 22M | (9) |

#### C.3 Training Hyperparameters

All models presented in this work were trained using an identical set of hyperparameters, eliminating the need for experiment- or sample-specific algorithmic tuning. The global loss function optimized by the network is computed as a simple, unweighted mean squared error (MSE) averaged across all four target outputs. Because the four target parameters $(D, N, B, \sigma_0)$ span multiple orders of magnitude (with $N$ and $B$ scaling in the thousands and $D$ and $\sigma_0$ scaling near 1) unweighted training on natural-space values leads to severe loss imbalance, where the network entirely ignores small-magnitude parameters during weight updates. To resolve this, log-transformation ensures the network prioritizes relative (multiplicative) errors over absolute additive errors. Concurrently, normalizing all four targets to a bounded $[0,1]$ range via min-max scaling prevents the large-magnitude parameters from entirely dominating the unweighted global MSE loss gradients as described in Equation 7.

Following standard ViT training protocols, network weights were optimized using the AdamW optimizer with a base learning rate of 0.001(11). In line with our regularization-heavy strategy, we applied a decoupled weight decay of 0.0001 alongside an attention and MLP dropout probability of 0.1 to prevent memorization of simulation-specific features. We trained the models for a total of 300 000 iterations and a batch size of 128. Crucially, because our simulation harness generates data continuously on-the-fly, every training sample is evaluated exactly once and never reused. In this infinite-data regime, the concept of a traditional 'training epoch' does not apply. We specifically choose to train for a similar duration as ImFCSNet (384 000 steps across 8 different learning rates in (2)) to ensure that any observed performance improvements are a result of architectural differences, rather than being due to extended training time.

To ensure training stability, we use a linear learning rate warmup over the first 10% of training (30 000 iterations) and use gradient clipping at a global norm of 2.0 to prevent exploding gradients (12, 13). Finally, Polyak weight averaging was applied with an exponential moving average (EMA) factor of 0.9999 to obtain final weights that are robust to step-wise batch noise/stochasticity (14).

Notably, our architecture is exceptionally lightweight in parameter count and requires minimal VRAM ($\sim$ 3.5 GB) during training, and $<$ 1 GB VRAM during inference, even for the comparatively heavier $3\times3$ ViT model. While the models evaluated in this work were trained on a GPU cluster (Nvidia V100) for iteration, we benchmarked the training pipeline on standard consumer hardware (Nvidia RTX 3060 Mobile, 6 GB VRAM; AMD Ryzen 7 5800H CPU, as used for runtime benchmarking). Continuous training for 300 000 steps on this consumer setup takes approximately 16 hours, confirming that customized model training and iteration remains possible even without datacentre hardware.

### D ALTERNATIVE CONFIGURATIONS

#### D.1 Single Pixel Models

In the main text, we introduced a single-pixel ($1\times1$) ImFCSViT variant to resolve sub-pixel structural heterogeneity in F-tractin datasets, relaxing the spatial homogeneity assumptions enforced by the canonical $3\times3$ model. While the core transformer blocks remain identical, adapting the ViT for optimal single-pixel inference requires a few changes:

1. We retain the simulation strategy detailed in Appendix C.1, but extract only the central pixel rather than the $3\times3$ region.
2. We reduce the maximum sampling limit for the diffusion coefficient in the data generation phase from 20.0 to 10.0 µm$^2$/s. This reflects the reduced observation area of a single pixel, which inherently limits the maximum observable molecular velocity before transit times drop below the frame rate.
3. We adjust the spatiotemporal tokenization from $3\times3\times10$ (90 pixels per token) to $1\times1\times20$ (20 pixels per token).
4. We proportionally reduce the embedding dimension (*i.e.*, the internal feature representation size) from 96 in the canonical ImFCSViT to 24 in the $1\times1$ variant.

This reduction in embedding size is the most critical architectural change. Given we parse the 2500-frame input into temporal tubelets of 20 frames ($1\times1\times20$), if we project the 20-dimensional tokens into the 96-dimensional embedding space used by the $3\times3$ model, we end up with an over-parameterized network, with an information expansion coefficient of $96\div20\approx4.8\times$. Empirically, this lack of a representational bottleneck allows the model to memorize the specific noise profiles of the simulations rather than learning generalized physical features. In practice, this over-parameterization causes severe regression to the mean, where the network outputs tend towards the average of the log-sampled training distribution rather than tracking the true kinetic signal. To prevent this, we proportionally reduce the embedding dimension of the $1\times1$ ImFCSViT to 24, enforcing an internal representation space that better maps to the information density present in each tokenized tubelet, allowing the network to learn dynamics that generalize across the simulation-to-experiment gap.

To highlight the performance consequences of removing spatial context, we evaluated the $1\times1$ ImFCSViT on the homogeneous DOPC bilayer preparations (Figure D.1). When mapped across a single 2500-frame window, the $1\times1$ spatial $D$-map (Figure D.1c) appears visually homogeneous, similar to the NLS $D$ map. Crucially, it entirely misses the highly hindered residual vesicles that were clearly identified by the $3\times3$ model. Consequently, its probability density is strictly unimodal (Figure D.1b), exhibiting a wider spread and a slight upward bias compared to the 50 000-frame NLS baseline. This inability

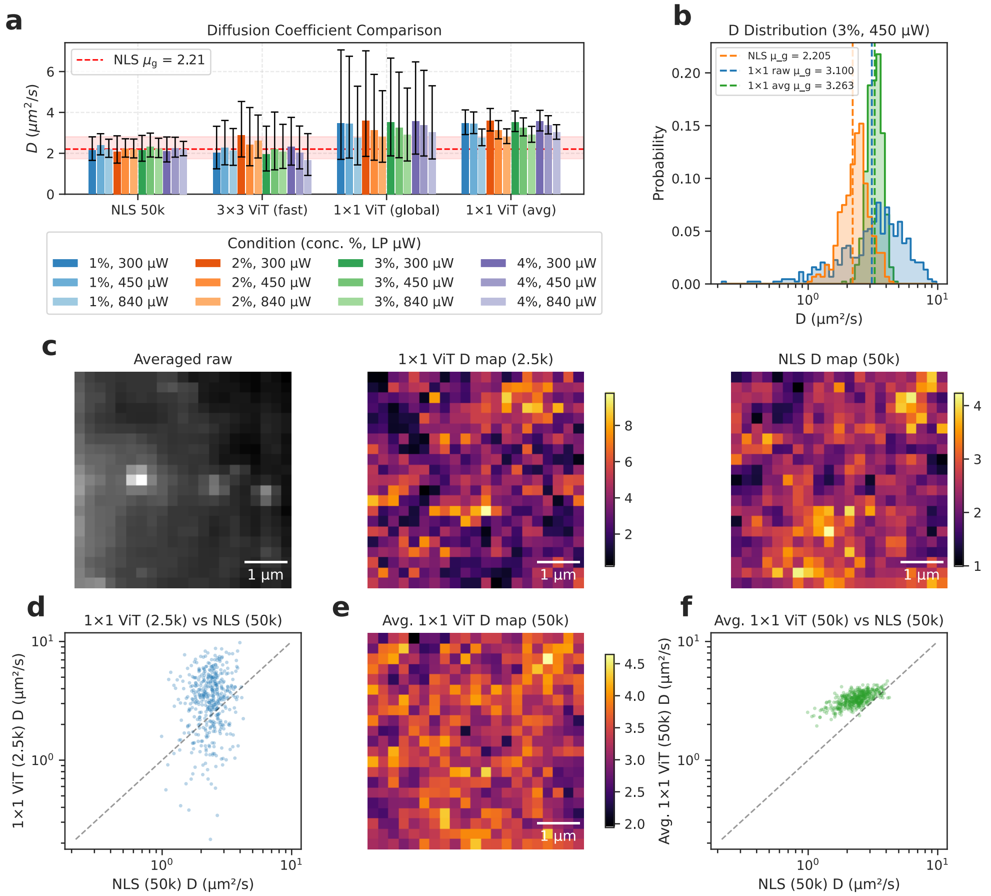


Figure D.1: Validation of $1 \times 1$ ImFCSViT diffusion analysis on DOPC supported lipid bilayers. (**a**) Diffusion coefficients ($D$) for NLS (control), 3×3 ViT (fast component only), 1×1 ViT (global, single 2.5k window), and 1×1 ViT (temporal average of all 20 prediction windows). Bars represent geometric mean ×/÷ geometric standard-deviation factor; the red band marks the geometric mean and SD interval of the pooled NLS measurements. (**b**) Log-scale distribution of $D$ for a representative measurement (3% concentration, 840 µW laser power), showing the unimodal 1×1 ViT distributions (single-window raw, light blue; temporal average, green) alongside the NLS reference (orange). Vertical lines indicate geometric means. (**c**) Temporally averaged 2500-frame region and the corresponding $1 \times 1$ ViT and NLS $D$ map, showing the inability of the $1 \times 1$ ViT to observe the slow-diffusing regions coinciding with high-intensity residual vesicles, similarly to NLS. (**d**) Scatter plot of per-pixel D values of the $1 \times 1$ ViT versus the NLS D values, showing that the ViT-derived diffusion coefficient at 2500 frames overpredicts compared to NLS-based fitting at 50 000 frames. (**e**) Temporally averaged $1 \times 1$ ViT map across all 20 available prediction frames, demonstrating the corresponding increase in precision, and ability to observe the structures. (**f**) Scatter plot of per-pixel D values of the temporally averaged $1 \times 1$ ViT across the full 50 000-frames and corresponding 20 predictions versus the NLS D values, showing that the ViT-derived diffusion coefficients correlates well with the NLS estimations at 50 000 frames post-temporal averaging.

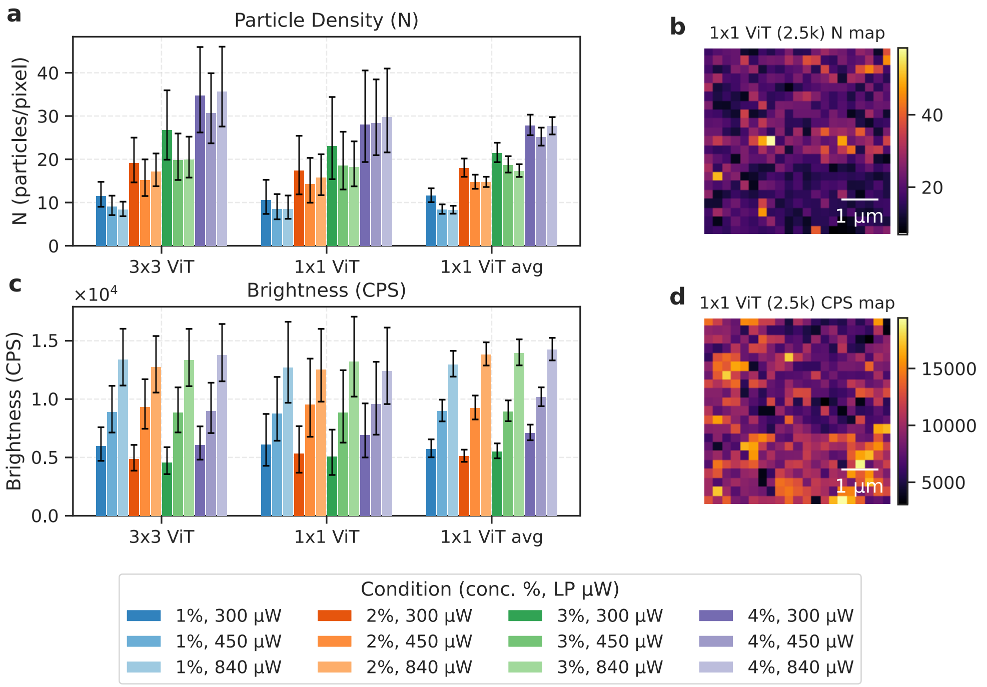


Figure D.2: Intensity-derived parameter estimates from the $1 \times 1$ ImFCSViT on DOPC supported lipid bilayers. (**a**) Particle density $N$ (per pixel) for the $3 \times 3$ ViT (global), $1 \times 1$ ViT (single 2.5k window), and $1 \times 1$ ViT after temporal averaging of all prediction windows. Bars represent geometric mean $\times/\div$ geometric standard-deviation factor for each model. (**b**) Representative $N$ map from the $1 \times 1$ ViT (single-window) for the 3% concentration, 450 µW condition. (**c**) Brightness (counts per second, CPS) barplots for the same three model variants; (**d**) Corresponding CPS map from the same representative measurement.

to observe slow components show that the spatial component of the $3 \times 3$ model is what allows it to isolate the fast- and slow-components, and the $1 \times 1$ model performs like NLS, being unable to distinguish hindered movement from background variance in a short one-dimensional intensity trace.

Finally, we replicate the temporal averaging strategy across all 20 prediction maps produced from the 50 000 frames, and visualize the overall $D$ map for the $1 \times 1$ model, and observe strong spatial correlations with the NLS standard over the same time window (Figure D.1e). The per-pixel scatter plot (Figure D.1f) shows the strong correlation with the NLS baseline across the same 50 000 window, just with a compressed dynamic range, proving that the network accurately extracts the underlying bulk dynamics once noise is averaged out.

We further investigated the $1 \times 1$ ImFCSViT using the particle density ($N$) and brightness (CPS) prediction heads. As shown in Figures D.2a and d, the $1 \times 1$ model manages to estimate similar $N$ and $CPS$ values as the $3 \times 3$ model despite using $9\times$ less data in terms of pixels processed. This delineates the difference in estimation difficulty between parameters: $D$ is a purely kinetic parameter heavily reliant on inter-frame fluctuations and spatial context, making it highly sensitive to shot noise. Conversely, $N$ and CPS are derived from the bulk statistical moments of the intensity trace (the mean and variance). These global trace statistics, aggregated efficiently via the normalization token, remain relatively stable even within a single pixel across 2500 frames.

Note that we deliberately choose to retain the point-spread-function prediction head ($\sigma_0$) in the single-pixel model. Inherently, a single pixel lacks the spatial structure to measure radial volume expansion. This was retained to enforce a strict network ablation study. Altering the number of target variables inherently reshapes the global MSE loss optimization landscape, re-weighting the backpropagation gradients across the shared components of the network. By forcing the $1 \times 1$ model to optimize for all four parameters exactly as its $3 \times 3$ counterpart, we ensured that spatial sequence sizing was the sole manipulated variable. The expected failure of the single-pixel model to accurately predict $\sigma_0$ is demonstrated explicitly using the induced defocusing dataset in Appendix A.1. This demonstrates the parameter crosstalk as expected from NLS-based fitting with Equation 2, where the parameters are inexplicably linked, further enforcing the strengths of the spatial component present in the $3 \times 3$ ViT over its single pixel counterpart.

#### D.2 Frame Counts

Throughout the main text, we design our ViT architecture to take in $3 \times 3$ ROIs, with a measurement time corresponding to 2500 frames. However, vision transformers do not enforce fixed input lengths, making it possible to extend or shorten the input frame lengths without any architectural changes. Unlike convolutional networks like ImFCSNet, where the flattened feature vector size is entirely dependent on the input dimensions, the ViT's attentive pooler aggregates any token sequence into a constant-sized vector.

To analyze the performance boundaries of our approach, we trained independent ViT models on 2000, 1500, 1000, 500, and 100 frames, and evaluate the DOPC bilayer samples used in the main text. Because of the ViT's architectural flexibility, these variants utilize the exact same code and structural hyperparameters as the canonical 2500-frame model; the network simply processes a dynamically shorter sequence (e.g., from 250 spatiotemporal tokens down to 10 tokens for the 100-frame case). However, we stress that each input length configuration is trained from scratch, as extrapolation to different input sizes (and thus sequence lengths) is not supported by our current use of fixed absolute positional embeddings.

In Figure D.3, we analyze the effect of different frame counts on the reference DOPC samples across differing concentrations and laser powers. Using the non-parametric Otsu method established previously, we isolate the fast-diffusing component and report its geometric mean and standard deviation across varying frame counts in Figure D.3a. As expected, lowering the frame count reduces the availability of statistics, resulting in a corresponding increase in the spread of estimated values. However, we find that even at 1000-frames, the $3 \times 3$ ImFCSViT is able to derive a fast-$D$ component that is consistent with the NLS-derived ground truth values. By then inspecting a selected representative sample (3% concentration, 450 µW), we observe that the model is clearly able to localize the slow component around the visible vesicles, even down to 500 frames, only failing catastrophically at 100 frames with $D$ maps that fail to properly isolate the immobile/slow-moving vesicles. Crucially, these estimates are achieved without temporal averaging, showing that we can go even lower than our proposed 2500-frame limit, with reliable estimates even at 1000 frames, a decrease of $50\times$ compared to the 50 000 frames required for reliable Imaging FCS parameter estimates.

To visualize the clear benefit of the increased temporal resolution, we then visualize the evolution of $D$ over time. In the case of conventional fitting, we only obtain a single parameter estimate per-pixel per-50 000 frames. For ImFCSViT, we instead get $\frac{50000}{\text{No. of frames}}$ estimates per-pixel per-50 000 frames. In Figure D.4, we can clearly see the effect of increased temporal resolution, going from 20 points in the 2500-frame case, up to 500 points in the 100-frame extreme. The same gradual decrease in precision can be observed, as well as the upwards bias that occurs at 500 input frames onwards (Figure D.4e-f). However, it is notable that ImFCSViT estimates remain consistent over time across all frame counts, and even down to 1000 frames (Figure

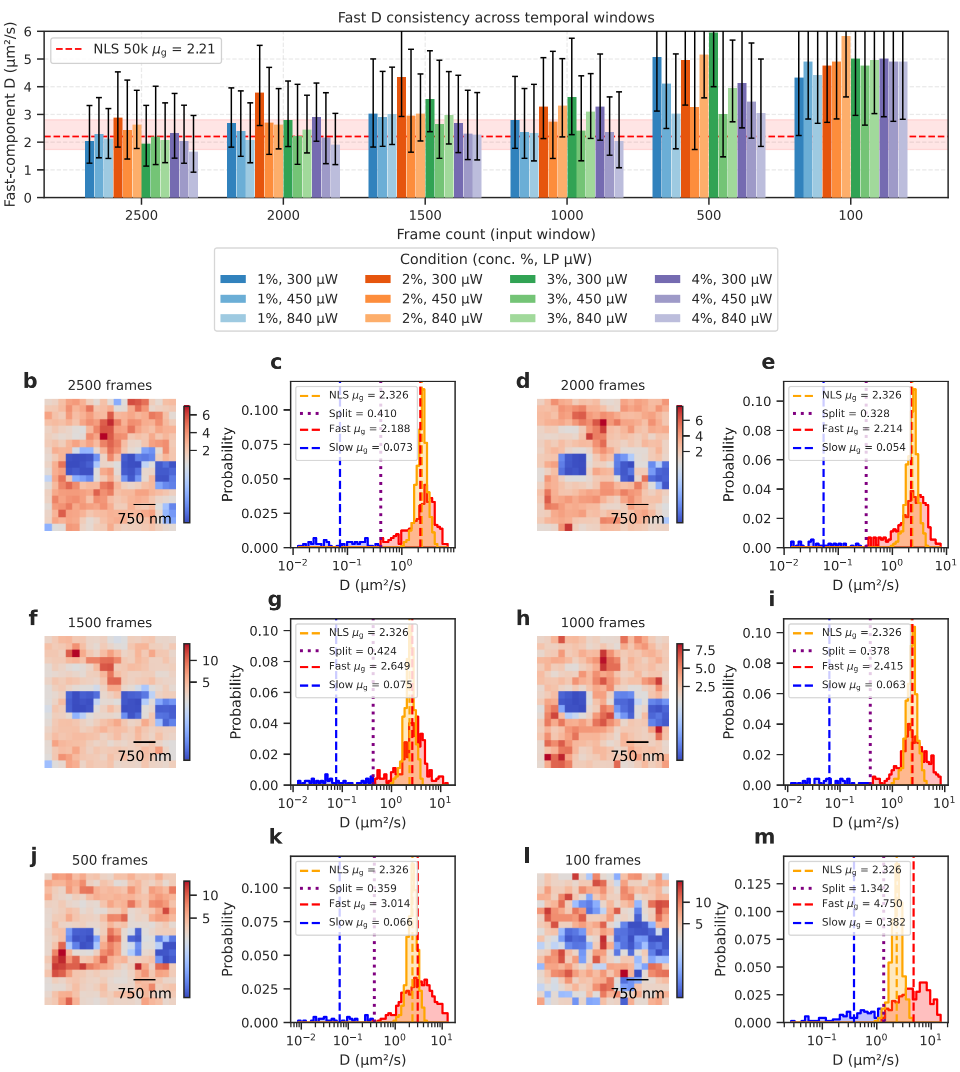


Figure D.3: Impact of input frame count on ImFCSViT diffusion predictions across DOPC supported lipid bilayers. **a** Fast-component $D$ values (geometric mean $\times/\div$ geometric standard-deviation factor) for all 12 experimental conditions (varying concentration and laser power) vs input window length. The red dashed line and shaded band denote the geometric mean and geometric SD interval of the conventional NLS fits (50 000 frames). All frame-count variants adopt identical transformer architecture and training hyperparameters; only the number of input frames. **b, d, f, h, j, l** Spatial $D$-maps for the selected representative measurement (3% concentration, 450 μW) at each frame count. Maps use a diverging colour scale centred on the per-model Otsu threshold; **c, e, g, i, k, m** Log-scale histograms overlying the NLS reference distribution and the ViT-D distributions at each frame count. As discussed in the Results section, bimodality is observed, with slow regions corresponding to residual vesicles. The Otsu-detected split point (purple) and corresponding fast (red) and slow (blue) populations with geometric means (dashed vertical lines) are marked.

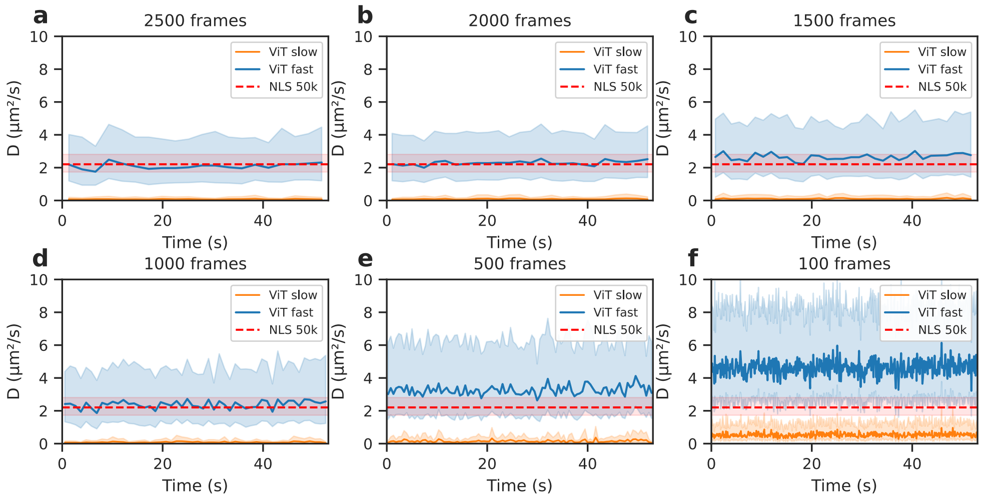


Figure D.4: Temporal evolution of fast and slow diffusion components across decreasing input window lengths for the representative DOPC sample (3% concentration, 450 µW). Each panel shows per-window geometric mean $D$ of the fast (blue) and slow (orange) components, separated by non-parametric Otsu thresholding as shown in Figure D.3, and the shaded band indicating the geometric standard-deviation factor (×/÷) interval. The red dashed line and band represent the global NLS reference (50 000 frames). As the window length shortens, we observe the spread of predictions increasing, and by 500 frames, there starts to be an observable upwards bias in the mean-$D$ value as well. However, the decrease in precision is made up for by the increased temporal resolution.

D.4d) demonstrate good agreement with NLS at 50 000 frames.

We investigate the other auxiliary outputs of our ImFCSViT model. The particle density $N$ predictions remain consistent down to 1000 frames (Figure D.5a), showing the relative ease of fitting a global descriptor in comparison to local descriptors like $D$ where frame-to-frame fluctuations are key. The brightness (CPS) predictor is stable down to 1500 frames (Figure D.5b). Finally, PSF $\sigma_0$ parameter shows relative consistency across all frame counts (Figure D.5c), though consistency does appear to suffer below 1500 frames.

Interestingly, we observe in both Figures D.3a and D.4f that the $D$ estimates at 100-frames are remarkably consistent despite the high upwards bias. The poor accuracy can be attributed to the extremely short 0.1 second measurement time, but this motivates a more in-depth exploration into whether this extreme configuration can be useful in situations where diffusion coefficient changes are required, but not necessarily quantitative values.

To this end, we apply these models on a set of 1,2-dimyristoyl-sn-glycero-3-phosphocholine (DMPC) bilayers acquired across multiple temperatures (preparation as described in Appendix B.1). DMPC has a known phase transition temperature of 24 °C, below which it exists in an ordered gel phase (hence a lower $D$), and above which it transforms to a disordered fluid phase (hence a higher $D$) (15, 16). We then apply the same slow- and fast-component splitting via Otsu thresholding across all available samples, and visualize the trend across all temperatures for the 2500-frame, 1000-frame, and 100-frame ImFCSViT variations (Figure D.6).

From Figure D.6a, we observe that the canonical 2500-frame ImFCSViT configuration is the best, effectively matching NLS across all temperatures and capturing the phase transition between 24 °C and 25 °C. With Figure D.6b, we find the 1000-frame configuration captures the same trend as NLS, but with a more drastic upwards bias. Finally, in the extreme case of Figure D.6c with only 100 frames, we find that ImFCSViT is no longer able to capture the minute differences across increasing temperatures, while also being subject to a large upwards bias. However, the phase transition between 24 °C and 25 °C is captured as an increase, hinting that the 100-frame configuration can be applied in cases where change detection, rather than exact quantification, is desired.

Finally, we note that one currently unexplored area is adapting models across different input lengths, or more specifically, training a model on one-length, but having it work on sequence lengths that it was not trained on. As our current model uses absolute sin/cos positional embeddings, the trained weights fail to extrapolate to different sequence lengths without finetuning (*i.e.*, retraining the model with the new frame count). However, this is an active area of research. Techniques such as Rotary Position Embeddings (RoPE) (17), which encode relative temporal distances, inherently handle variable input lengths, and can be adopted into fluorescence bioimaging-focused ViTs if true frame count-agnostic ViTs are required.

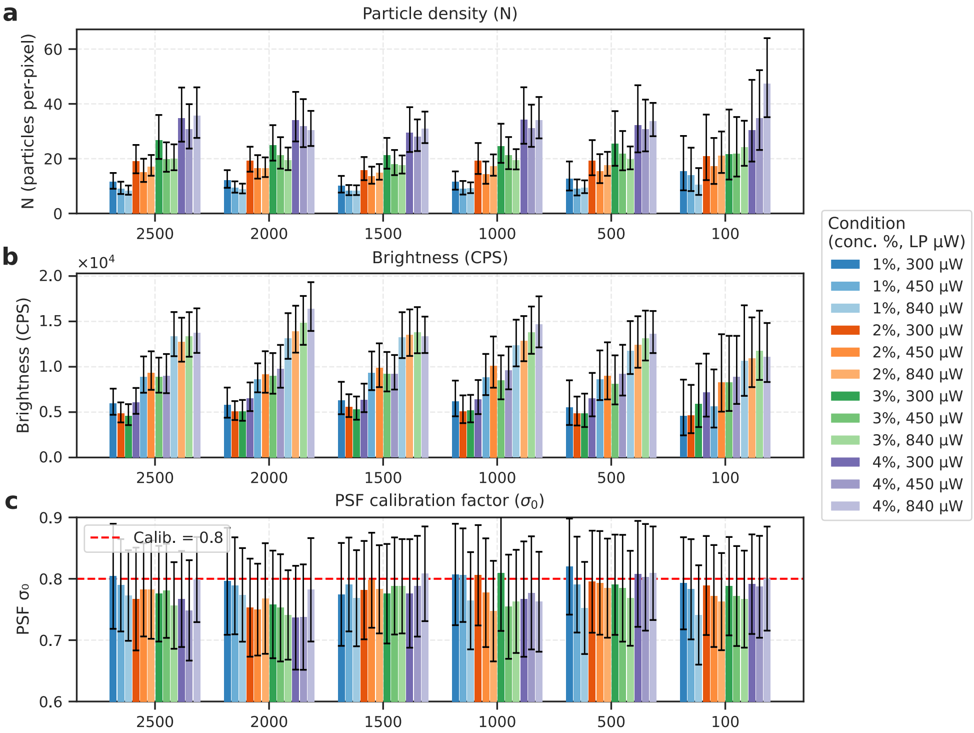


Figure D.5: Stability of intensity-derived parameters across input window lengths for DOPC supported lipid bilayers. (**a**) Particle density $N$ estimated by the $3 \times 3$ ImFCSViT. Geometric mean and geometric standard-deviation factor are shown for each condition (12 combinations of label concentration and laser power); bars are grouped by concentration first to highlight the invariance with laser power. (**b**) Brightness (counts per second, CPS) from the same model. Conditions are sorted by laser power first to reveal the proportional scaling with illumination intensity. (**c**) PSF calibration factor $\sigma_0$ (arithmetic mean $\pm$ SD) across frame counts; the red dashed line marks the calibrated reference value of 0.8. Error bars for $N$ and CPS span $\times/\div$ one geometric standard-deviation factor; PSF errors are $\pm 1$ standard deviation.

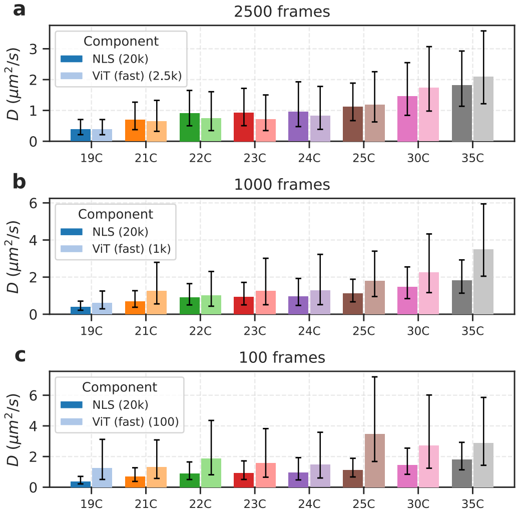


Figure D.6: Aggregated diffusion coefficients for DMPC bilayers across different temperatures. (**a**) ViT fast-$D$ estimate obtained using only 2500 frames, evaluated with the 2500-frame ViT, as indicated by lighter bars on the right. Reference NLS-based $D$ estimates with 20 000 frames are provided, as indicated by darker bars on the left. (**b**) Same as **a**, but evaluated with the 1000-frame ViT. (**c**) Same as **a**, but evaluated with the 100-frame ViT. Error bars span $\times/\div$ one geometric standard-deviation factor.

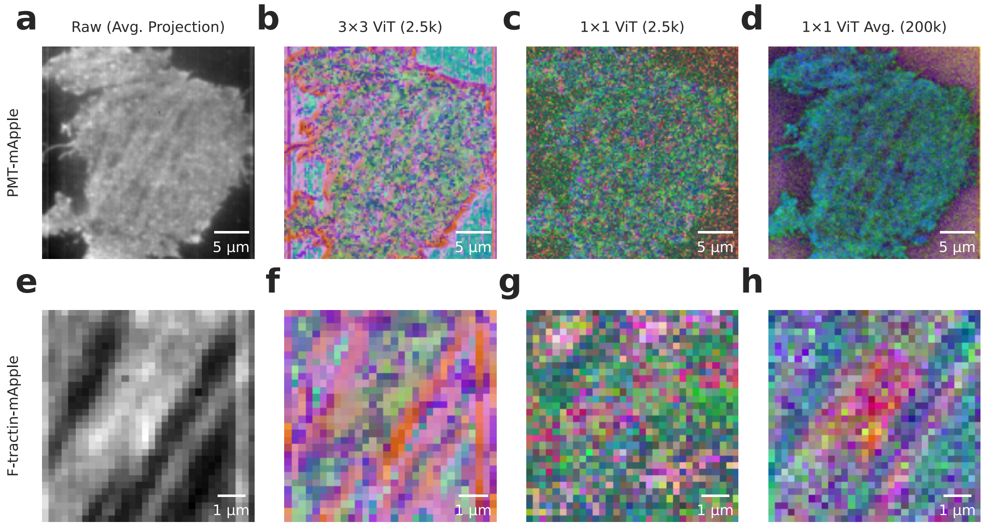


Figure E.1: Visualization of the learned latent feature spaces in the $3 \times 3$ and $1 \times 1$ ImFCSViT models. High-dimensional feature embeddings were extracted, reduced to three dimensions via Principal Component Analysis (PCA), and mapped to RGB channels for spatial visualization. (**a**–**d**) Feature representations for the PMT-mApple dataset. (**a**) Raw fluorescence image, time-averaged over the first 2,500 frames. (**b**) PCA-projected features from the $3 \times 3$ ViT at the initial frame chunk ($t = 0$), demonstrating robust spatial awareness. (**c**) Corresponding single-frame features from the $1 \times 1$ ViT, showing high spatial noise due to independent pixel-wise processing. (**d**) Temporal average of the $1 \times 1$ ViT features prior to PCA projection, showing that structure can still be extracted once averaged over time. (**e**–**h**) Analogous feature visualizations for the F-tractin-mApple sample containing actin filaments, confirming the failure case of the $3 \times 3$ model, and recovery of fine fibrous structures in the $1 \times 1$ model through temporal averaging.

### E INTERPRETABILITY

As deep learning-based neural networks are generally considered black boxes, we attempt to examine the internal feature representations of our ViT architecture. More specifically, we bypass the linear regression layer of our network (Stage 3 in Figure 1), visualizing and interpreting the model's internal representations to potentially explain why the network works and fails on certain tasks.

#### E.1 Network Features

First, we examine the network's pre-regression features. Specifically, we extract the length-96 feature vectors produced by the attentive pooler prior to parameter estimation. To obtain a rich, diverse feature space, we evaluate the uncropped PMT-mApple CHO-K1 cell measurement from Figure 4, which contains a complex mixture of on-cell membrane regions, cell boundaries, and background pixels.

As interpreting features in a high-dimensional space is not feasible visually, we use an approach inspired by (18), and use principal component analysis (PCA) for dimensionality reduction, transforming the 96-dimensional space into 3-dimensions. We stress our use of PCA, which is a linear dimensionality reduction technique, as opposed to non-linear methods like UMAP or t-SNE, which inherently induce structure and artificial clustering. This ensures that any observed feature similarities represent true linear correlations in the network's latent space rather than clustering-induced artifacts. To then interpret these principal components, we map the 3 PCA components to RGB values. This effectively gives a similarity mapping of different $3 \times 3 \times 2500$ regions, where regions that produce similar features are assigned similar colours.

As shown in Figures E.1 b and E.1 f, we observe the $3 \times 3$ ViT's rich feature space on the cell, producing a feature map with the similar structural pattens observed in the diffusion coefficient map from Figure 4b, corresponding the identified bimodality of fast (green) and slow (pink) diffusing regions. However, we also observe the ViT's susceptibility to cell edges, where the network features are significantly different from the successful on-cell fits. This is a similar effect to the failures of the $3 \times 3$ ViT on the F-tractin data with spatial non-homogeneity, as edge pixels correspond to $3 \times 3$ regions that include a mixture of on-cell and off-cell regions, which is a case that the model has not been trained on, and is thus out-of-distribution. On the F-tractin-mApple cell visualizations (Figure E.1f), we observe the network's noisy feature space that is affected by the spatial

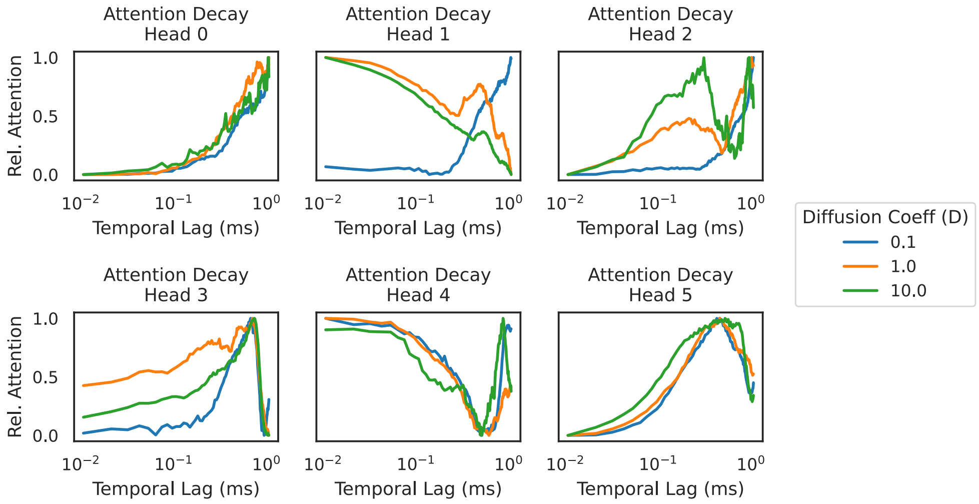


Figure E.2: Visualization of the attention decay for each of the 6 attention heads in the $3 \times 3$ ViT, coloured by the simulated diffusion coefficient. Notably, each head exhibits a distinct attention decay profile, with Head 4 exhibiting a temporal attention decay shape similar to a classical ACF.

non-homogeneity of the actin filament-rich environment, only producing features that are consistent (green) on the regions where homogeneity is possible across a $3 \times 3$ space.

Extending our analysis further to the $1 \times 1$ ViT, we repeat the same process, the only difference being the input size (single pixels instead of $3 \times 3$ regions) and the feature vector lengths (24 instead of 96). When evaluating just the first 2500 frames, the resulting feature maps are visibly dominated by noisy features (Figures E.1c and E.1g). While we can observe some semblance of on-cell and off-cell separation for the PMT-mApple cell (Figure E.1c), noise dominates even in feature space. For the F-tractin-mApple sample, structural morphology is entirely lost (Figure E.1g).

In the Results section, we demonstrated that the high variance of the single-window $1 \times 1$ ViT predictions could be resolved by temporally averaging the parameter estimates over 80 independent sequential windows (Figure 5j-l). By temporally integrating these feature vectors before applying PCA, we integrate out the stochastic shot noise (Figures E.1d and E.1h). The resulting temporally-averaged $1 \times 1$ feature maps recover the underlying cellular structure to a high degree of fidelity, visually confirming that the model learns representations that map to true physical morphology once trace-level variance is suppressed.

#### E.2 ViT Attention Visualization

Next, we examine the self-attention patterns of the $3 \times 3$ ViT. We probe this across simulations with diffusion coefficients spanning two orders of magnitude ($D = 0.1\mu m^2/s$, $1.0\mu m^2/s$, and $10.0\mu m^2/s$). Inspired by classical ACF analysis, we evaluate the network's attention weights as a function of the temporal distance between tokens. Because each token encodes 10 frames, examining token lags $\tau$ allows us to map the network's internal temporal dependencies across discrete 10-frame intervals.

Specifically, excluding the global normalization token, the self-attention mechanism in each transformer attention head maps the relationship between all 250 sequential tokens (each representing a $3 \times 3 \times 10$ chunk), producing a 2-dimensional attention matrix of size $250 \times 250$. We then analyze the attention weights as a function of the temporal lag $\tau$ between tokens to understand how these temporal dependencies contribute to the ViT's final feature representations. As transformer-based attention is bidirectional, we average both the forward $(+\tau)$ and backward $(-\tau)$ token relationships to compute the mean attention at a given token lag $\tau$. By averaging the attention weights of all token pairs separated by a specific lag $\tau$, we effectively perform a ViT-specific analogue of the time-averaging process used to compute classical ACFs. To exclude attention lags with low statistics (*i.e.*, long token lags that correspond to a low number of samples), we truncate our token lags to a maximum lag of 100 tokens. Similarly, we exclude lag-0 attention, as this architectural self-similarity dominates the attention matrix and obscures the inter-token temporal dynamics we wish to analyze (analogous to ACF-based data fitting excluding lag-0 due to the dominance of shot-noise). Finally, to obtain smooth decay profiles, we ensemble-average these lag-dependent attention weights across the initial 2,500-frame chunks for 50 independent simulations per $D$ value, apply min-max normalization independently per head, and visualize the decay on a logarithmic temporal axis.

From Figure E.2, we observe that all 6 attention heads show different attention decay patterns. Perhaps most surprisingly, head 4 learns a pattern that is resembles the shape of an ACF, even representing the expected leftwards shift of the ACF as the diffusion coefficient increases. Head 1 also shows an attention decay pattern that appears to be diffusion dependent (high long-term attention with slow diffusion, high short-term attention with fast diffusion). The rest of the heads could correspond to features that contribute to the other predictors ($N$, CPS, $\sigma_0$), but a thorough analysis of each specific head is beyond the scope of this paper.

While the spontaneous emergence of ACF-like attention patterns strongly supports the hypothesis that the network is deriving physical representations from raw data, these visualizations should be treated as qualitative heuristics. In deep learning, attention weights do not equate to strict mathematical feature importance, and interpreting individual heads is inherently ad-hoc. Rather, we highlight that the ViT architecture provides built-in mechanisms for structural interpretability, giving us another dimension to probe network fitting decisions that is inaccessible to convolutional architectures. Whether the ViT works should be based primarily on the fitting results, which we have established throughout this paper.

### F ABLATIONS

To validate specific architectural decisions made in the design of our ViT-based framework, we performed a series of ablation studies. Specifically, we compare against the convolutional ImFCSNet baseline (2), and we demonstrate the necessity of our proposed Normalization Token for simultaneous multi-parameter prediction.

To ensure a fair comparison, all ablation models (including ImFCSNet variants) were trained from scratch using the exact same simulated dataset, loss normalization bounds, and optimization hyperparameters (e.g., learning rate schedules, weight decay) as our canonical ViT model. All models share the same $3 \times 3 \times 2500$ input size.

We train the following configurations, comparing them against the canonical $3 \times 3$ ImFCSViT

1. $3 \times 3$ **ViT (no norm)**: Standard ImFCSViT training, but with no normalization token or Z-score normalization step. Highlights the significance of the normalization token as an informative prior.

2. **ImFCSNet (D)**: Reproduction of ImFCSNet, only predicting $D$, and with input Z-score normalization. We use the same simulation settings and training schedules from Appendix C, showing that our regularization-heavy settings are not ViT-specific, and work well for convolutional architectures as well.

3. **ImFCSNet (all)**: ImFCSNet, modified for 4-target prediction ($D$, $N$, CPS and $\sigma_0$). As discussed in the Vision Transformers section of the Materials and Methods section, input normalization removes the statistics necessary for $N$ and brightness estimation, so we remove Z-score normalization.

To evaluate all models and ablations, we analyze the performance of each setting on the DOPC SLB dataset. For the case of $D$, we observe that 3 of the 4 models are able to separate the fast and slow components (Figure F.1b,d,f), the exception being **ImFCSNet (all)**, which fails to produce a bimodal distribution (Figure F.1h-i). While the $3 \times 3$ ViT is able to identify the existence of slow-moving residual vesicles, the fast-$D$ estimates are less accurate compared to the canonical $3 \times 3$ ViT, indicating the importance of normalization for $D$ fitting (Figure F.1a). Similarly to the canonical $3 \times 3$ ViT, **ImFCSNet (D)** with input normalization is able to capture the right diffusion coefficients, and in some cases (concentration of 4%) able to be more accurate than the $3 \times 3$ ViT. Perhaps most importantly, these ablations further enforce the fact that the ability to identify slow-moving residual vesicles is not a ViT-specific property, but rather granted by the spatial component of the $3 \times 3$ models.

For the other targets, we exclude **ImFCSNet (D)** as it lacks $N$, CPS and $\sigma_0$ predictors. From Figure F.2, we find that the canonical $3 \times 3$ ImFCSViT performs the best in terms of consistency. The modified 4-target ImFCSNet (all) fails to estimate parameters consistently (with the exception of $\sigma_0$). Similarly, the $3 \times 3$ **ViT (no norm)** also fails to achieve the consistency of the canonical model for $N$ and CPS (Figure F.2a,b). This shared drop in performance demonstrates that the failure is not strictly architectural, but rather stems from the fundamentally conflicting requirements of the target variables. Tracking localized frame-to-frame fluctuations (for $D$) requires suppressing bulk global statistics, while extracting particle density (for $N$ and CPS) requires them. This suggests that the normalization token itself is a strong informative prior that acts as an early-fusion global context vector. Without it, the network's representational capacity is overloaded by attempting to simultaneously deduce bulk global statistics and track localized frame-to-frame fluctuations. By providing this prior explicitly, we design the network with a built-in way of separating these parameters. This is further hinted at by the $3 \times 3$ **ViT (no norm)**'s inability to fit PSF (Figure F.2c).

These ablations help give us confidence that our proposed normalization token modification to the standard ViT is principled, and an optimal way to maintain strong fitting capabilities across both dynamics and intensity-derived fit parameters.

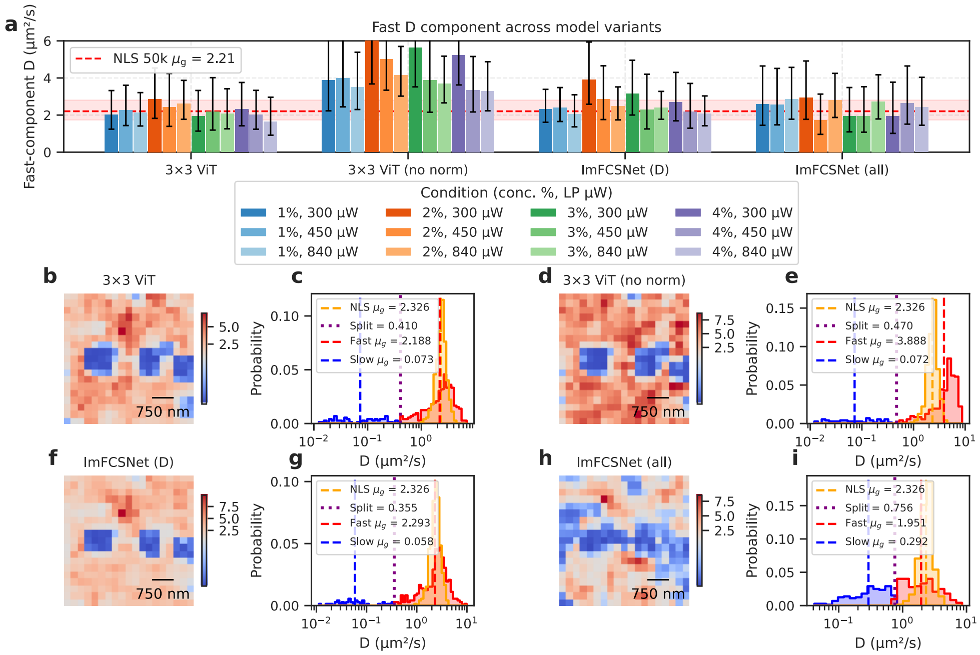


Figure F.1: Diffusion predictions for different $3 \times 3$ models and ablations across DOPC supported lipid bilayers. **a** Fast-component $D$ values (geometric mean $\times/\div$ geometric standard-deviation factor) for all 12 experimental conditions (varying concentration and laser power) for baseline $3 \times 3$ ViT model against ablations. The red dashed line and shaded band denote the geometric mean and geometric SD interval of the conventional NLS fits (50 000 frames). **b, d, f, h** Spatial $D$-maps for the selected representative measurement (3% concentration, 450 µW) for each ablation. Maps use a diverging colour scale centred on the per-model Otsu threshold; **c, e, g, i** Log-scale histograms overlying the NLS reference distribution and the ViT or ImFCSNet $D$ distribution. As discussed in the Results section, bimodality is observed, with slow regions corresponding to residual vesicles. The Otsu-detected split point (purple) and corresponding fast (red) and slow (blue) populations with geometric means (dashed vertical lines) are marked.

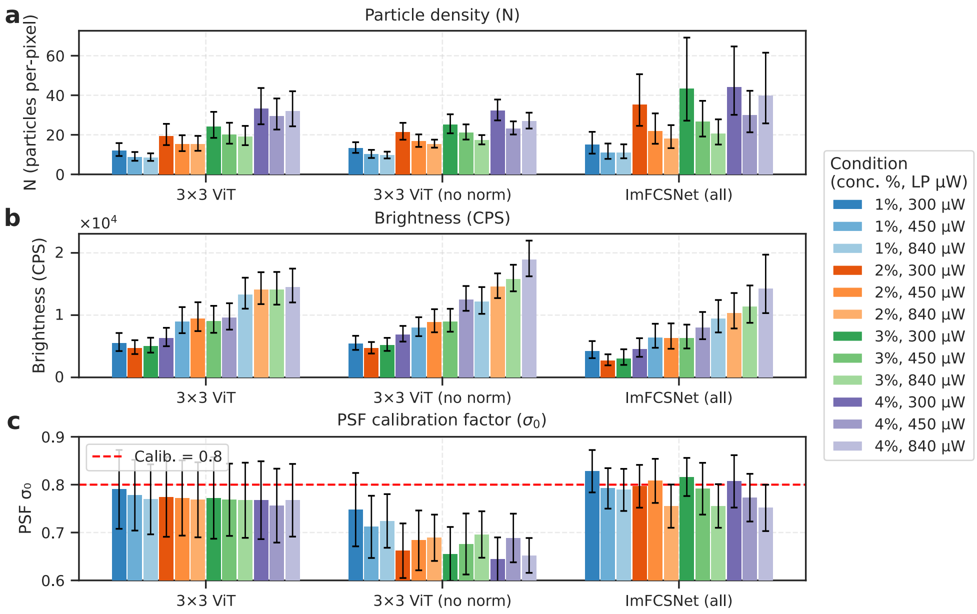


Figure F.2: Stability of intensity-derived parameters for different $3 \times 3$ models and ablations for DOPC supported lipid bilayers. (**a**) Particle density $N$ estimated by the $3 \times 3$ ViT. Geometric mean and geometric standard-deviation factor are shown for each condition (12 combinations of label concentration and laser power); bars are grouped by concentration first to highlight the invariance with laser power. (**b**) Brightness (counts per second, CPS) from the same models. Conditions are sorted by laser power first to reveal the proportional scaling with illumination intensity. (**c**) PSF calibration factor $\sigma_0$ (arithmetic mean $\pm$ SD) across frame counts; the red dashed line marks the calibrated reference value of 0.8. Error bars for $N$ and CPS span $\times/\div$ one geometric standard-deviation factor; PSF errors are $\pm 1$ standard deviation.

### REFERENCES


1. Nirmalya Bag et al. "Calibration and Limits of Camera-Based Fluorescence Correlation Spectroscopy: A Supported Lipid Bilayer Study". In: *Chemphyschem : a European journal of chemical physics and physical chemistry* 13.11 (2012), pp. 2784–2794. ISSN: 1439-7641. DOI: 10.1002/cphc.201200032.
2. Wai Hoh Tang et al. "Deep Learning Reduces Data Requirements and Allows Real-Time Measurements in Imaging FCS". In: *Biophysical Journal* 123.6 (2024), pp. 655–666. ISSN: 0006-3495. DOI: 10.1016/j.bpj.2023.11.3403.
3. Harikrushnan Balasubramanian et al. "The Dependence of EGFR Oligomerization on Environment and Structure: A Camera-Based N&B Study". In: *Biophysical Journal* 121.23 (2022), pp. 4452–4466. ISSN: 0006-3495. DOI: 10.1016/j.bpj.2022.11.003.
4. L. K. Tamm and H. M. McConnell. "Supported Phospholipid Bilayers". In: *Biophysical Journal* 47.1 (1985), pp. 105–113. ISSN: 0006-3495. DOI: 10.1016/S0006-3495(85)83882-0.
5. Shambhavi Pandey, Nithin Pathoor, and Thorsten Wohland. "Super-Resolution Algorithms for Imaging FCS Enhancement: A Comparative Study". In: *Biophysical Journal* 124.20 (2025), pp. 3428–3440. ISSN: 0006-3495, 1542-0086. DOI: 10.1016/j.bpj.2025.03.031.
6. Shambhavi Pandey and Thorsten Wohland. "EGFR Does Not Directly Interact with Cortical Actin: A SRRF'n'TIRF Study". In: *Biophysical Journal* 123.21 (2024), pp. 3736–3749. ISSN: 0006-3495. DOI: 10.1016/j.bpj.2024.09.022.
7. Thorsten Wohland et al. "FCS Videos: Fluorescence Correlation Spectroscopy in Space and Time". In: *Biochimica et Biophysica Acta (BBA) - General Subjects* 1868.11 (2024), p. 130716. ISSN: 0304-4165. DOI: 10.1016/j.bbagen.2024.130716.
8. Michael Hirsch et al. "A Stochastic Model for Electron Multiplication Charge-Coupled Devices–from Theory to Practice". In: *PloS One* 8.1 (2013), e53671. ISSN: 1932-6203. DOI: 10.1371/journal.pone.0053671.
9. Alexey Dosovitskiy et al. *An Image Is Worth 16x16 Words: Transformers for Image Recognition at Scale*. 2021. DOI: 10.48550/arXiv.2010.11929. arXiv: 2010.11929 (cs).
10. Hugo Touvron et al. *Training Data-Efficient Image Transformers & Distillation through Attention*. 2021. DOI: 10.48550/arXiv.2012.12877. arXiv: 2012.12877 (cs).
11. Ilya Loshchilov and Frank Hutter. "Decoupled Weight Decay Regularization". In: *arXiv preprint arXiv:1711.05101* (2017). arXiv: 1711.05101.
12. Razvan Pascanu, Tomas Mikolov, and Yoshua Bengio. *On the Difficulty of Training Recurrent Neural Networks*. 2013. DOI: 10.48550/arXiv.1211.5063. arXiv: 1211.5063 (cs).
13. Andreas Steiner et al. *How to Train Your ViT? Data, Augmentation, and Regularization in Vision Transformers*. 2022. DOI: 10.48550/arXiv.2106.10270. arXiv: 2106.10270 (cs).
14. B. T. Polyak et al. "Acceleration of Stochastic Approximation by Averaging". In: *SIAM Journal on Control and Optimization* 30.4 (1992), pp. 838–855. DOI: 10.1137/0330046.
15. Wen Chen et al. "Determination of the Main Phase Transition Temperature of Phospholipids by Nanoplasmonic Sensing". In: *Scientific Reports* 8.1 (2018), p. 14815. ISSN: 2045-2322. DOI: 10.1038/s41598-018-33107-5.
16. D. Needham and E. Evans. "Structure and Mechanical Properties of Giant Lipid (DMPC) Vesicle Bilayers from 20.Degree.C below to 10.Degree.C above the Liquid Crystal-Crystalline Phase Transition at 24.Degree.C". In: *Biochemistry* 27.21 (2002), pp. 8261–8269. ISSN: 0006-2960. DOI: 10.1021/bi00421a041.
17. Jianlin Su et al. *RoFormer: Enhanced Transformer with Rotary Position Embedding*. 2023. DOI: 10.48550/arXiv.2104.09864. arXiv: 2104.09864 (cs).
18. Maxime Oquab et al. *DINOv2: Learning Robust Visual Features without Supervision*. 2024. DOI: 10.48550/arXiv.2304.07193. arXiv: 2304.07193 (cs).
